# Intra-species quantification reveals differences in activity and sleep levels in the yellow fever mosquito, *Aedes aegypti*

**DOI:** 10.1101/2024.03.16.585223

**Authors:** Oluwaseun M. Ajayi, Emily E. Susanto, Lyn Wang, Jasmine Kennedy, Arturo Ledezma, Angeli’c Harris, Evan S. Smith, Souvik Chakraborty, Nicole E. Wynne, Massamba Sylla, Jewelna Akorli, Sampson Otoo, Noah H. Rose, Clément Vinauger, Joshua B. Benoit

## Abstract

*Aedes aegypti* is an important mosquito vector of human disease with a wide distribution across the globe. Climatic conditions and ecological pressure drive differences in the biology of several populations of this mosquito, including blood-feeding behavior and vector competence. However, no study has compared activity and/or sleep among different populations/lineages of *Ae. aegypti*. Having recently established sleep-like states in three mosquito species with observable differences in timing and amount of sleep among species, we investigated differences in activity and sleep levels among 17 *Ae. aegypti* lines drawn from both its native range in Africa and its invasive range across the global tropics. Activity monitoring indicates that all the lines show consistent diurnal activity, but significant differences in activity level, sleep amount, number of sleep bouts, and bout duration were observed among the lines. Variations in specific activity and sleep parameters were explained by differences in host preference, ancestry, and human population density for the lineages collected in Africa. This study provides evidence that the diurnal sleep and activity profiles for *Ae. aegypti* are consistent, but there are significant population differences for *Ae. aegypti* sleep and activity levels and interactions with humans may significantly impact mosquito activity and sleep.

## Introduction

The yellow fever mosquito, *Aedes aegypti*, is the primary vector of several arboviruses, and an epidemiologically important species with a wide distribution, especially in tropical and subtropical areas (Eisen and Moore, 2013; Brown et al., 2014; Kraemer et al., 2015). A strong relationship exists between *Ae. aegypti* and humans based on a suite of observed ecological behaviors including indoor dwelling and breeding, endophilic resting, multiple anthropophilic, and daytime blood-feedings (Scott et al., 2000; Scott and Takken, 2012; Li et al., 2017). This mosquito species plays an essential role in the emergence and expansion of arboviral diseases, including dengue fever, Chikungunya, and Zika (Nuckols et al., 2015; Weaver and Lecuit, 2015; Cauchemez et al., 2016). Due to the worldwide distribution of *Ae. aegypti*, environmental differences drive subtle differences in the biology of these mosquitoes from different geographical zones. Differences in geographical origins and lineages influence specific characteristics such as vector competence, blood versus sugar feeding, and host preferences (Ponlawat and Harrington, 2005; Spencer et al., 2005; Olson et al., 2020; Rose et al., 2020). Nonetheless, the influence of lineage differences on other aspects of mosquito biology, especially daily/circadian rhythms in *Ae. aegypti* is yet to be evaluated.

Like other insects, mosquitoes inhabit environments that experience daily oscillations in environmental factors such as light and temperature across a typical 24-hour period (Rund et al., 2016; Brody, 2020). In response to the daily cycling of environmental light, mosquitoes have evolved to display time-dependent rhythms of a large proportion of their biology, including sugar feeding (Yee and Foster, 1992), olfaction (Rund et al., 2013), oviposition (Fritz et al., 2008; Chadee, 2010), mating (Benelli, 2015), metabolism (Gray and Bradley, 2003), immunity (Murdock et al., 2013), and activity patterns (Jones et al., 1972; Peterson, 1980; Kawada and Takagi, 2004). Apart from the tight connection between feeding behavior and locomotor activity, feeding in mosquitoes and other blood-feeding insects is rhythmic and can be used to predict when infectious bites will occur (Schlein and Warburg 1986, Yee and Foster 1992, Lorenzo and Lazzari 1998, Fritz et al. 2014). Moreover, the light/dark cycle also drives mosquito interaction with other organisms, including human hosts, ultimately influencing pathogen transmission (Rund et al., 2016). Broad-scale differences in the temporal organization of activity are well known between different mosquito genera, where field observations have established increased night-time (nocturnal) and daytime (diurnal) biting in *Anopheles* spp and *Aedes* spp, respectively (Tuchinda et al., 1969; Rund et al., 2016). Additionally, studies in the *An. gambiae* and *Culex pipiens* complexes revealed that activity patterns and levels vary based on differences in strain (Shinkawa et al., 1994; Rund et al., 2012). Still, these studies only examined two or three lineages.

Along with daily/circadian rhythms, there has been a recent interest in understanding sleep behavior in mosquitoes and its epidemiological importance. Although investigations are still in their infancy, targeted studies using mosquitoes of the genera *Aedes*, *Culex,* and *Anopheles* have shown that sleep-like states are characterized by stereotyped postures, increased arousal threshold, consolidated immobility in specific periods of the circadian day, and sleep recovery after prior sleep deprivation, consistent with what has been reported in other insects (Helfrich-Förster, 2018; Ajayi et al., 2020; Ajayi et al., 2022; Ajayi et al., 2023). Sleep deprivation in *Ae. aegypti* results in reduced host landing and blood-feeding propensity, suggesting that sleep might play an important role in mosquito pathogen transmission (Ajayi et al., 2022). This study also shows that circadian timing and duration of sleep differ among mosquito species, matching what is known about day and night active species (Veronesi et al., 2012; Rund et al., 2016; Rund et al., 2020; Ajayi et al., 2022). However, unlike in mosquitoes, sleep phenotypes in insects can differ even among populations/lineages of the same species as shown in several studies in *Drosophila melanogaster* (Harbison et al., 2013; Harbison et al., 2017; Serrano Negron et al., 2018). Hence, evaluating if similar observations are observable in mosquitoes is important.

In this present study, we quantified the differences in activity and sleep levels across multiple lines/populations of *Ae. aegypti* from West Africa (Rose et al., 2020) and established lab strains, which encompassed 17 mosquito populations. Our results indicate the commonality of diurnal behavior, but significant differences in activity levels, sleep amount, sleep bout number, and bout duration occurred among different lines. Differences in host preference, ancestry, and human population density based on previously established phenotypes (Rose et al., 2020) significantly influenced the variation in some activity and sleep metrics in the West African populations. A second paper on population differences among *Ae. albopictus* has been developed in conjunction with this study and reveals differences among populations for that species (Wynne et al., 2024).

## Materials and methods

### Mosquito husbandry

Seventeen *Aedes aegypti* lines were used to examine activity and sleep profiles. Seven of these lines have a West African origin, namely; PK10, Kedougou, Kintampo, Kumasi, Mindin, Ngoye, and Thies (Rose et al., 2020). The other ten lines *i.e.*, Black Eyes (Liverpool), Costa Rica, D2MEB, Gainesville (default laboratory strain at the University of Cincinnati which serves as the control), Indiana, Liverpool, Orlando, Puerto Ibo, Puerto Rico, and Rockefeller have been established across different laboratories in the United States. The West African lines were provided by collaborators at Princeton University (Princeton, NJ, USA) and in their countries of origin, while the others were acquired from BEI Resources (Manassas, VA, USA). Colonies of all the lines were established at the University of Cincinnati, with at least two gonotrophic cycles completed before being used for experiments. Moreover, the Liverpool and Rockefeller lines were also reared and established at Virginia Tech for comparison studies across different sites.

All colonies were maintained at 25°C, 80% relative humidity (RH) under a 15 h:9 h light: dark (L/D) cycle with access to water and 10% sucrose *ad libitum*. Eggs were produced from 2- to 4-week-old females through blood feeding with a human host (University of Cincinnati IRB 2021-0971). After egg hatching, larvae were separated into 18×25×5 cm containers (at a density of 250 individuals per container) and were fed finely ground fish food (Tetramin, Melle, Germany) and maintained in an incubator at 24°C, 70–75% RH, under a 12 h:12 h L/D cycle until adult emergence. Emerged adult mosquitoes had access to water and 10% sucrose *ad libitum*. All adult female mosquitoes used for the experiment were 7–14 days post-ecdysis. As the experimenters represent potential hosts to the mosquitoes, activity, and sleep assays were conducted in an enclosed room in an isolated building that was not accessed during experiments to eliminate potential disturbances.

### Activity and sleep quantification

The activity and sleep profiles and levels of the multiple lines of *Ae. aegypti* were quantified using a Locomotor Activity Monitor 25 (LAM25) system developed for *Drosophila* (TriKinetics Inc., Waltham, MA, USA) and adapted for mosquitoes and ticks (Rund et al., 2012; Lima-Camara et al., 2014; Rosendale et al., 2019; Ajayi et al., 2022; Benoit et al., 2023). For each line, individual mosquitoes were loaded into 25×150 mm clear glass tubes with access to water and 10% sucrose provided *ad libitum*. Tubes were then placed horizontally in the LAM25 system, which consists of 32 monitor channels in an ‘8×4’ horizontal by vertical matrix, allowing the simultaneous recording of 32 mosquitoes during a single trial. Multiple lines were monitored during each trial by loading individual mosquitoes randomly into the channels to prevent bias. The system was placed in an incubator at 24°C, 70–75% RH, under a 12 h:12 h L/D cycle (11 h full light, 11 h full darkness, and one h dawn and one h dusk transitions).

Activity level was calculated as the number of infrared beam breaks made by the mosquito in a minute. Based on an earlier study, sleep was quantified using a period of inactivity (lack of beam breaks) lasting 120 minutes (Ajayi et al., 2022). Each line was measured until at least a sample size of 40 individuals was achieved. The mosquitoes were measured for seven days, excluding two days of acclimation. Data was analyzed using the Rethomics framework in R with associated packages such as *behavr*, *ggeth*o, *damr*, and *sleep*r (Geissmann et al., 2019). For the Gainsville line, data collected from a previous study was used in the analyses (Ajayi et al., 2022). Mosquitoes that survived until the end of the recording were included in activity and sleep assessment. Three individuals with the least and three with the highest number of beam breaks were excluded from the analysis for lines with sample sizes less than 50 (representing 5% of 50 to the nearest whole number) to remove potential outliers. Lines with sample sizes greater than 50 but less than 100 had four individuals excluded (representing 8% of 50). In addition, ten individuals on both ends were marked as outliers for the Liverpool and Rockefeller lines assayed at Virginia Tech (representing 10% of 100, which matches the outlier removal for samples from UC).

### Data analysis

Experimental replicates used in this study are distinct samples and biologically independent. Sample sizes are reported in the figure legends. The following response variables were computed from data analyzed using Rethomics: total activity level, daytime activity level, nighttime activity level, total sleep amount, daytime sleep amount, nighttime sleep amount, sleep bout number, daytime bout duration, nighttime bout duration, and peak periodicity for both activity and sleep. Linear models were fitted as appropriate to determine the influence of line differences on these variables, where “line” represents the explanatory variable for all the models in question. To ascertain statistical significance for each model, *F*-tests using the “anova” function were computed where *p* < 0.05 was considered statistically significant (Chambers and Hastie, 1992). Post-hoc comparisons among the lines were made using *emmeans* (default - Tukey method), where *p* < 0.05 was also considered statistically significant (Searle et al., 1980). Post-hoc comparisons for all models are shown in the supplemental tables.

In addition, data on host preference index, ancestry, and human population density obtained from another study (Rose et al., 2020), were included as explanatory variables in single-variable and combined-variables linear mixed models (where “line” was used as a random effect) for only the West African populations. The Akaike information criterion (AIC) values were used to select the model with the better fit between the single-variable and combined-variables model for each response variable (Chakrabarti and Ghosh, 2011). The host preference indices of these West African populations range from -0.58 (depicting a strong preference for non-human hosts) to 0.69 (showing a strong preference for human hosts) (Rose et al., 2020). Based on genomic data, the differences in phenotypic traits among these West African lines are attributed to differences in ancestry components related to the two major subspecies, *Ae. aegypti aegypti* (Aaa) and *Ae. aegypti formosus* (Rose et al., 2020; Rose et al., 2023), where increasing Aaa abundance is known to be associated with urban environments. All analyses were done in R v. 4.2.1, where the “lm” function was used for the linear models (Wilkinson and Rogers, 1973), “emmeans” for post-hoc comparisons, and “lmer” was used for the linear mixed models (Kuznetsova et al., 2017).

## Results

### Multiple lines show diurnal behavior in activity with increased sleep in the night

Diurnality of activity is consistent across all the lines assayed in this experiment (See Figures S1A - S17A). For most of the lines, activity increased from mid-day till the onset of the scotophase (dark phase), and there was a reduction of activity in the scotophase relative to the photophase (light phase). However, some West African lines, including Thies, PK10, Mindin, and Kedougou, showed higher activity levels during specific periods of the night. It is also noteworthy to state that some *Ae. aegypti* lines, *i.e.,* Rockefeller, Puerto-Ibo, Liverpool, Indiana, D2MEB, and Costa Rica, showed strong activity peaks as they approached the start of sunrise. As expected, since sleep is an inverse of activity, the lines all showed decreased sleep during the photophase and increased sleep during the scotophase (Figure S1B - S17B). The summary of averages and statistical outputs are in Tables S1 - S12.

### Differences in lines explain variations in activity levels and sleep amounts

We evaluated the differences in activity levels (beam counts) and amount of sleep among all the populations. Total average beam counts (both day and night, Figure 1A), daytime, and nighttime average beam counts (Figures 1B and 1C) were all significantly influenced by the line identity (total, *F*_16,697_ = 19.583, *p* < 0.001; daytime, *F*_16,697_ = 39.700, *p* < 0.001; nighttime, *F*_16,697_ = 5.425, *p* < 0.001). The greatest variation in beam counts due to line differences was observed during the daytime alone (adjusted R^2^ = 0.465) than in the whole day (adjusted R^2^ = 0.294) and nighttime alone (adjusted R^2^ = 0.090). The same scenario was seen with the amount of sleep among the multiple populations for total (both day and night, Figure 2A), daytime and nighttime (Figures 2B and 2C), where they were significantly influenced by line identity (total, *F*_16,697_ = 53.770, *p* < 0.001; daytime, *F*_16,697_ = 59.790, *p* < 0.001; nighttime, *F*_16,697_ = 15.770, *p* < 0.001). As expected, the most extensive variation in sleep amount due to line differences was seen in the daytime alone (adjusted R^2^ = 0.5688) in comparison to the whole day (adjusted R^2^ = 0.5422) and nighttime alone (adjusted R^2^ = 0.2490). As predicted, the mean daily activity and sleep amount showed a strong negative correlation (Spearman’s rank correlation = −0.8652, *p* < 0.001), which means that lines with high activity have short sleep duration and lines with low activity have long sleep duration.

**Figure 1:**
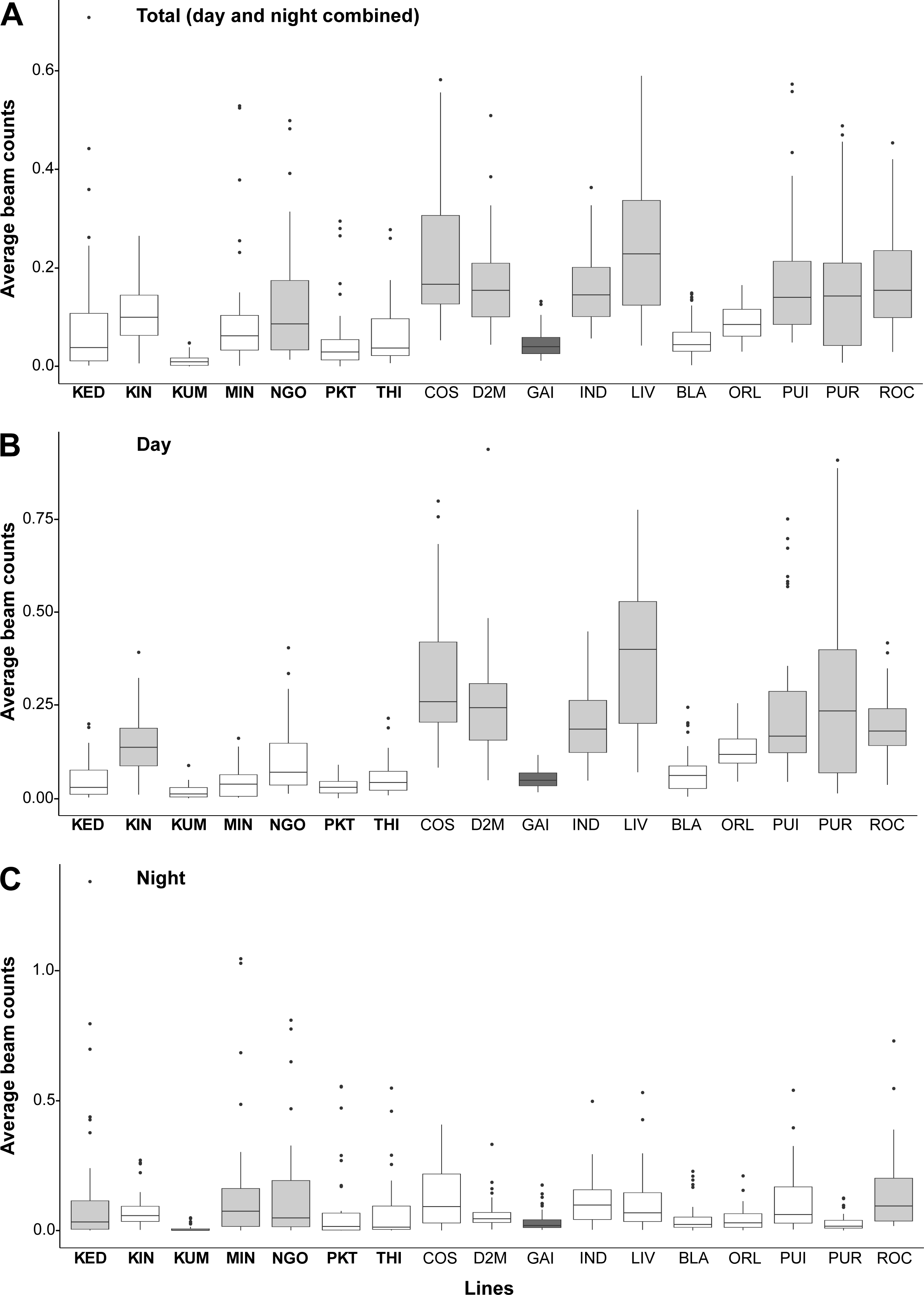
Activity levels differ across multiple *Aedes aegypti* lines. Comparison of average beam counts during (A) day and night, (B) daytime, and (C) nighttime among the different *Ae. aegypti* lines. KED: Kedougou (n=40); KIN: Kintampo (n=43); KUM: Kumasi (n=45); MIN: Mindin (n=41); NGO: Ngoye (n=36); PKT: PK10 (n=37); THI: Thies (n=34); COS: Costarica (n=40); D2M: D2MEB (n=37); GAI: Gainsville (n=52); IND: Indiana (n=48); LIV: Liverpool (n=36); BLA: Liverpool (Black eyes) (n=54); ORL: Orlando (n=48); PUI: Puerto Ibo (n=41); PUR: Puerto Rico (n=37); ROC: Rockfeller (n=45). Box plots represent the lower 25th percentiles, medians, and upper 75th percentiles. The labels in bold fonts (x-axis) represent the West African populations. The lines represented by the light grey-colored box plots are significantly different (*p* < 0.05) from the Gainsville line (dark grey-colored box plot) based on linear models used to assess significant differences in average beam counts among the lines. Estimated marginal means and pairwise post-hoc comparisons for all the lines are shown in Tables S1 - S6.

**Figure 2:**
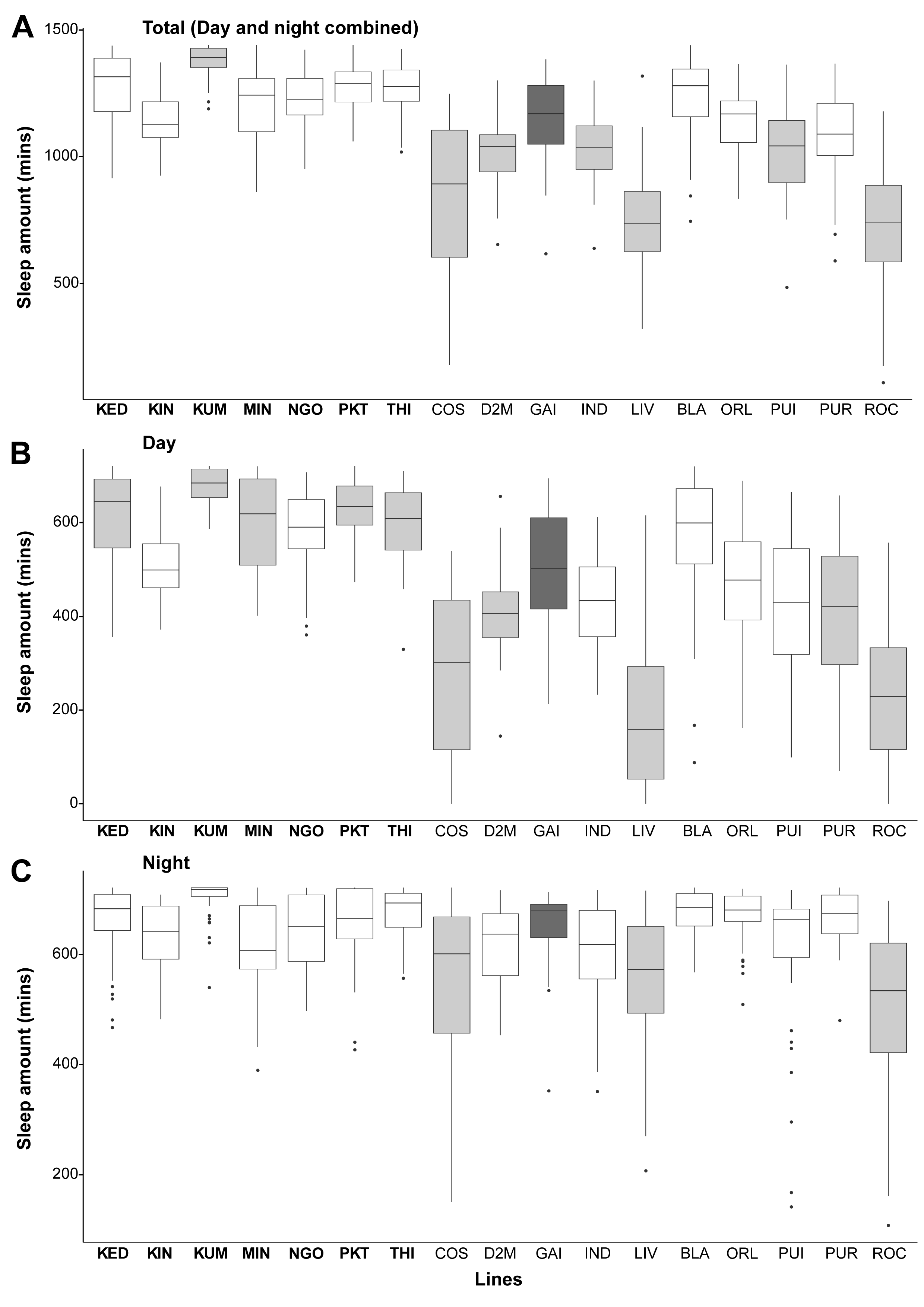
The duration of sleep differs across multiple *Aedes aegypti* lines. Comparison of average sleep amount during (A) both day and night, (B) daytime only, and (C) nighttime only among the different *Ae. aegypti* lines. KED: Kedougou (n=40); KIN: Kintampo (n=43); KUM: Kumasi (n=45); MIN: Mindin (n=41); NGO: Ngoye (n=36); PKT: PK10 (n=37); THI: Thies (n=34); COS: Costarica (n=40); D2M: D2MEB (n=37); GAI: Gainsville (n=52); IND: Indiana (n=48); LIV: Liverpool (n=36); BLA: Liverpool (Black eyes) (n=54); ORL: Orlando (n=48); PUI: Puerto Ibo (n=41); PUR: Puerto Rico (n=37); ROC: Rockfeller (n=45). Box plots represent the lower 25th percentiles, medians, and upper 75th percentiles. The labels in bold fonts (x-axis) represent the West African populations. The lines represented by the light grey-colored box plots are significantly different (*p* < 0.05) from the Gainsville line (dark grey-colored box plot) based on linear models used to assess significant differences in sleep amount among the lines. Estimated marginal means and pairwise post-hoc comparisons for all the lines are shown in Tables S7 - S12.

### Variation in other aspects of sleep architecture is influenced by differences in lines, but peak periodicity does not differ

We evaluated the impact of lines on other aspects of sleep architecture, including bout number and average bout duration across five (5) days and peak periodicity of both activity and sleep. Differences in line significantly impact variability in sleep bout number across multiple days (Figure 3A, *F*_16,697_ = 22.983, *p* < 0.001), explaining about 33% of the variation (adjusted R^2^ = 0.3303). This result was also seen in average bout duration, where differences in line significantly influence observed variability (Figure 3B, *F*_16,697_ = 15.002, *p* < 0.001). The percentage of individuals showing significant peak periodicity of locomotor activity and sleep differs for each line (**Table 1**). However, there are no significant differences in the peak periodicity of both locomotor activity (*F*_16,451_ = 1.274, *p* = 0.2097) and sleep (*F*_16,513_ = 1.049, *p* = 0.4029) across the multiple populations, where peak periodicity is approximately 24hrs.

**Figure 3:**
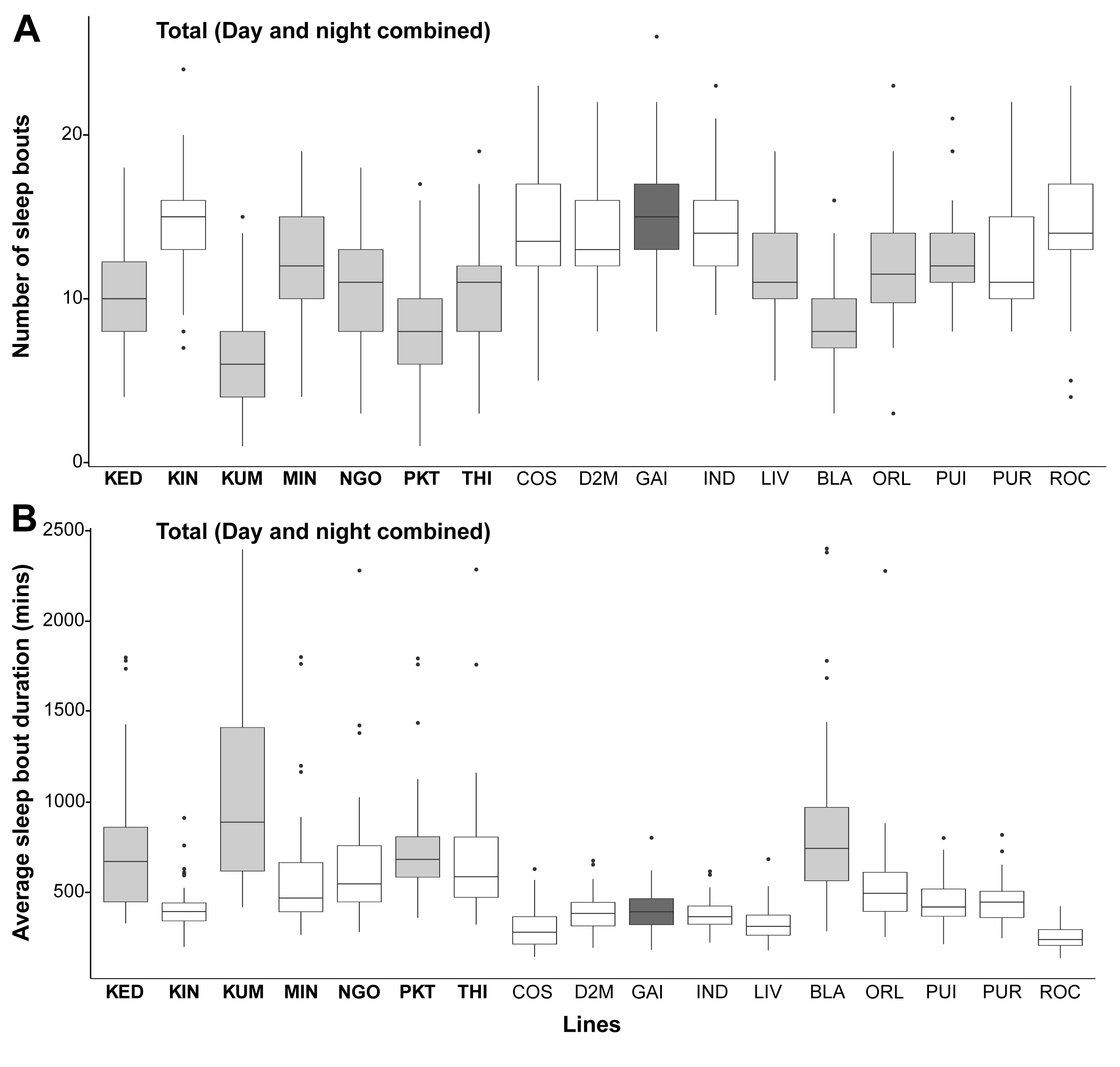
Number of sleep bouts and average bout duration differ across multiple *Aedes aegypti* lines. Comparison of (A) the number of sleep bouts, and (B) average sleep bout duration across five (5) days among the different *Ae. aegypti* lines. KED: Kedougou (n=40); KIN: Kintampo (n=43); KUM: Kumasi (n=45); MIN: Mindin (n=41); NGO: Ngoye (n=36); PKT: PK10 (n=37); THI: Thies (n=34); COS: Costarica (n=40); D2M: D2MEB (n=37); GAI: Gainsville (n=52); IND: Indiana (n=48); LIV: Liverpool (n=36); BLA: Liverpool (Black eyes) (n=54); ORL: Orlando (n=48); PUI: Puerto Ibo (n=41); PUR: Puerto Rico (n=37); ROC: Rockfeller (n=45). Box plots represent the lower 25th percentiles, medians, and upper 75th percentiles. The labels in bold fonts (x-axis) represent the West African populations. The lines represented by the light grey-colored box plots are significantly different (*p* < 0.05) from the Gainsville line (dark grey-colored box plot) based on linear models used to assess significant differences in the number of sleep bouts and average sleep bout duration among the lines. Estimated marginal means and pairwise post-hoc comparisons for all the lines are shown in Tables S13 - S16. Note: In (B), individuals with only a single bout of sleep across the five days were removed for graphical purposes.

**Table 1:** The number and percentage of individuals showing peak periodicity of activity and sleep for each measured line.

| Line | Sample size | No. of individuals showing peak periodicity (%) |  |
| --- | --- | --- | --- |
|  |  | Locomotor activity | Sleep |
| Kedougou | 40 | 22 (55.0) | 21 (52.5) |
| Kintampo | 43 | 38 (88.4) | 38 (88.4) |
| Kumasi | 45 | 21 (46.7) | 20 (44.4) |
| Mindin | 41 | 20 (48.8) | 26 (63.4) |
| Ngoye | 36 | 10 (27.8) | 19 (52.8) |
| PK10 | 37 | 15 (40.5) | 14 (37.8) |
| Thies | 34 | 9 (26.5) | 19 (55.9) |
| Costa Rica | 40 | 36 (90.0) | 37 (92.5) |
| D2MEB | 37 | 35 (94.6) | 37 (100) |
| Gainsville | 52 | 27 (51.9) | 32 (61.5) |
| Indiana | 48 | 34 (70.8) | 44 (91.7) |
| Liverpool | 36 | 33 (91.7) | 35 (97.2) |
| Liverpool (BLK) | 54 | 33 (61.1) | 38 (70.4) |
| Orlando | 48 | 37 (77.1) | 37 (77.1) |
| Puerto Ibo | 41 | 39 (95.1) | 40 (97.6) |
| Puerto Rico | 37 | 30 (81.1) | 32 (86.5) |
| Rockefeller | 45 | 29 (64.4) | 41 (91.1) |

BLK: Black eyes

### Strain difference in sleep and activity is consistent across sites

As most variation in activity and sleep is seen during the day periods, we compared the difference of day activity and sleep between two lines measured at the labs of both the University of Cincinnati (UC) and Virginia Tech (VT) to confirm if differences observed between lines are similar when tested at different locations. The relative difference of day beam counts between the Liverpool and Rockefeller lines (*i.e.* LVP/Rock) was not significantly different across sites (*W* = 1089, *p* = 0.7721), where the Liverpool line is approximately 1.97 and 2.05 fold higher than the Rockefeller line at the UC and VT sites, respectively. This is the same for day sleep, where the relative difference between the two lines is consistent across the sites (*W* = 905, *p* = 0.2771). For day sleep, the Liverpool line is approximately 0.74 and 0.85 fold lower than the Rockefeller line at the UC and VT sites, respectively. This suggests that sleep and activity variations between lines are repeatable when assessed by different experimenters at different locations.

### Host preference and other ecological variables show an association with activity and sleep parameters among the West African lines

We quantified the association of host preference, ancestry, and human population density with some activity and sleep parameter differences among the West African *Ae. aegypti* populations using linear mixed models for both single-variable and combined-variables analyses. According to genomic data in previous studies, these lines show variation in the proportion of *Aaa*-like ancestry, which influences host preference and other traits (Rose et al., 2020; Rose et al., 2023). Based on AIC, the combined-variables scenarios were used to examine the interaction between activity and sleep parameters with line-specific traits because of lower values for all the explanatory variables (except for only one instance) compared with the single-variable comparisons (**Tables 2A & B**).

**Table 2A:** The influence of host preference, ancestry, and human population density on some activity and sleep parameters (single-variable mixed models with line as a random effect).

| Model | Summary | Response variables tested |  |  |  |
| --- | --- | --- | --- | --- | --- |
|  |  | Day activity | Night activity | Day sleep | Night sleep |
| ~Preference | *p* | 0.666 | 0.663 | 0.987 | 0.611 |
|  | df | 7.080 | 7.584 | 7.144 | 7.285 |
|  | AIC | -764.392 | -159.510 | 3219.174 | 3110.733 |
| ~Ancestry | *p* | 0.362 | 0.576 | 0.714 | 0.745 |
|  | df | 7.103 | 7.661 | 7.162 | 7.293 |
|  | AIC | -765.082 | -159.642 | 3219.031 | 3110.897 |
| ~hpd | *p* | 0.674 | 0.119 | 0.321 | 0.067 |
|  | df | 6.999 | 6.700 | 7.012 | 6.961 |
|  | AIC | -764.381 | -161.871 | 3218.121 | 3107.452 |

hpd: Human population density; df: Degrees of freedom; AIC: Akaike information criterion. $p < 0.05$ indicates the significance of the explanatory variable with respect to the response variable in each fitted model. All values are in the nearest 3 decimal places. The df residual is 272 for all models. The explanatory variables of interest are defined previously (Rose et al., 2020).

Our models show a significant relationship between host preference and variation in day activity among our West African populations (**Table 2B)**. There is an association of ancestry with day activity and day sleep, and variation in night activity and night sleep is influenced by human population density among the tested populations (**Table 2B**). Interestingly, there is an increase in day activity for populations with a higher preference for non-human hosts. Also, there is a general decrease in night activity and an increase in night sleep with lines collected from urban areas with higher human density. Nevertheless, these correlations are not statistically significant and have weak coefficients.

**Table 2B:** The influence of host preference, ancestry, and human population density on some activity and sleep parameters (combined-variables mixed models with line as a random effect).

| Model | Variable of interest | Summary | Response variables tested |  |  |  |
| --- | --- | --- | --- | --- | --- | --- |
|  |  |  | Day activity | Night activity | Day sleep | Night sleep |
| ~preference+<br>ancestry+<br>hpd | Preference | *p* | 0.015 | 0.563 | 0.079 | 0.303 |
|  |  | df | 7.123 | 7.829 | 7.083 | 7.300 |
|  | Ancestry | *p* | 0.007 | 0.243 | 0.047 | 0.235 |
|  |  | df | 7.139 | 7.923 | 7.098 | 7.331 |
|  | hpd | *p* | 0.107 | 0.014 | 0.084 | 0.041 |
|  |  | df | 6.892 | 6.461 | 6.851 | 6.834 |
|  | AIC |  | -768.627 | -162.345 | 3217.581 | 3109.776 |

hpd: Human population density; df: Degrees of freedom; AIC: Akaike information criterion. $p < 0.05$ indicates the significance of the explanatory variable with respect to the response variable in each fitted model. Cells with significant $p$ values are shaded in grey. All values are in the nearest 3 decimal places. The df residual is 270 for all models. The explanatory variables of interest are defined previously (Rose et al., 2020).

## Discussion

In this study, we examined the differences in activity and sleep among multiple lines of the yellow fever mosquito, *Ae. aegypti*. Diurnal activity is consistent in all lines, but differences were seen among the lines for total activity, daytime activity, nighttime activity, total sleep, daytime sleep, nighttime sleep, number of sleep bouts, and duration of sleep bout. Our investigation shows that peak periodicity for both activity and sleep did not differ among the different lines, and differences in daytime activity and sleep between two representative lines were consistent when measured independently at two locations. In addition, depending on the models of choice, host preference, ancestry, and human population density influenced the differences observed in specific activity and sleep variables among our West African lines.

In a previous study, we observed diurnal activity in a single strain of *Ae. aegypti* mosquito (Ajayi et al., 2022), but the result from this current study, where an additional 16 populations were assayed, reinforces support for a widespread occurrence of diurnality in *Ae. aegypti*. Similar results of diurnal behavior were also observed in earlier studies where measurements were done using the Rockefeller and West African strains (Taylor and Jones, 1969; Lima-Camara et al., 2014; Rivas et al., 2018; Araujo et al., 2020; Teles-de-Freitas et al., 2020). Our current study is one of several studies where strain differences did not impact chronotype. Quantification in *An. gambiae* mosquitoes reported that the M and S forms (now considered two distinct species) maintained consistent daily nocturnal rhythms of flight activity under 12 h: 12 h L/D cycle conditions (Rund et al., 2012). Under a 16 h: 8 h L/D photoperiod, all strains (except one) of *Cx. pipiens molestus* from different geographical regions also showed similar entrained activity patterns (Shinkawa et al., 1994), similar to what was observed between lines in this current study. In *Cx. annulirostris*, geographically isolated populations monitored using video techniques all displayed crepuscular/nocturnal locomotor activity patterns (Williams and Kokkinn, 2005). Apart from mosquitoes, eight (8) field lines of the male Linden Bug, *Pyrrhocoris apterus*, with different free-running periods exhibited similar locomotor activity patterns, with one activity peak during the light phase under 18 h: 6 h L/D cycle (Pivarciova et al., 2016). Most insect species examined to this point have a specific time when peak activity occurs during each daily cycle.

Although we observed consistent diurnal behavior in our multiple lines, the activity and sleep levels differ considerably (Figure S1-S17). We observed differences in activity level (average number of beam crosses) and sleep amount (period of inactivity lasting at least 120 minutes) among the different lines, which were more pronounced during the photophase (daytime). More muted differences were noted between some lines during the scotophase (nighttime). Similar differences in activity levels between lines or populations have been documented in previous studies, even though the daily cycles are relatively similar. The activity levels are remarkably different between the field lines of *P. apterus* (Pivarciova et al., 2016), which can vary up to 100%. Irrespective of sex, a significant difference in daily total flight activity was observed between the M and S forms of *An. gambiae*, and Iran populations of *Cx. pipiens molestus* were more active than the Japanese populations, even though each species had the same activity profile (Shinkawa et al., 1994; Rund et al., 2012). In fruit flies, specific studies have linked geographical and genetic differences to differences in locomotor activity and sleep parameters. Nighttime levels of activity differ substantially between higher- and lower-latitude populations for *D. melanogaster*, and increased sleep correlates with proximity to the equator (Svetec et al., 2015). Genome-wide association (GWA) studies in *D. melanogaste*r showed variable sleep amount and waking activity among genetically diverse DFRP and SIP lines, and even among flies with identical genotypes (Harbison et al., 2013; Harbison et al., 2017; Serrano Negron et al., 2018). It is important to state that it was not reported if these selected lines exhibit similar activity/sleep patterns.

Sleep architecture (sleep bouts and average bout duration) differs across the different lines in this study, where some lines display more sleep fragmentation than others, like the Gainsville, Kintampo, and Rockefeller lines. These sleep differences are not surprising as geographically or genetically different populations of selected lines of *D. melanogaster* display differences in other sleep traits alongside variation in sleep amount (Harbison et al., 2013; Svetec et al., 2015; Harbison et al., 2017; Brown et al., 2018; Serrano Negron et al., 2018). Specifically, a study of *D. melanogaster* populations from North and Central America demonstrated that sleep bouts of equatorial populations are approximately twice as long as the temperate populations, suggesting that the former populations prefer less-fragmented sleep (Svetec et al., 2015).

Free-running period (FRP; close to 24 h), which is used to confirm the occurrence of circadian rhythm in insects, might be influenced by strain differences. In the *Cx. pipiens* complex, geographical differences influence the length of the free-running activity period in which the Iran strain has a significantly shorter period than the other strains (Shinkawa et al., 1994). In a similar study on *P. apterus*, FRPs varied remarkably among the 59 field lines originating from different localities, with the slowest clock at approximately 28 h (Pivarciova et al., 2016). Contrary to these observations, the males of both molecular forms of *An. gambiae* showed shorter periods than females, but there was no observable strain-specific difference in the FRP length of activity (Rund et al., 2012). This result was also seen in *Drosophila*, where there was no significant difference in the FRPs between the Panama City and Maine populations (Svetec et al., 2015). The difference between these studies and the other investigations is that comparisons of FRPs were only done between two strains in the *Anopheles* and *Drosophila* studies. We have no data for free-running periods since we could not quantify activity-rest rhythms under constant darkness. However, we can show that peak periodicity of activity and sleep (under L/D condition) does not vary significantly among our populations. This result is unsurprising as the populations all followed the direct influence of the light *Zeitgeber*. Since these two indices (FRP and peak periodicity) might not be directly comparable, it is important for future studies on *Ae. aegypti* to compare FRPs among strains, even if it is for a few different populations.

It has been established that host choice in mosquitoes is primarily influenced by host odor (Takken and Verhulst, 2013), where *Ae. aegypti* from human-specialist and generalist populations strongly prefer the odors of humans and non-human animals, respectively (Gouck, 1972; McBride et al., 2014). Odor preference between humans and animals of several African lines (including those used in our study) varied significantly among lines, in which Thies and Ngoye showed a clear human preference, and others either showed no preference or preferred animal hosts (Rose et al., 2020). These lines show significant variation in ancestry components, especially the proportion of *Aaa*-like ancestry, which influences traits such as host preference, suggesting the importance of these traits in the successful exploitation of humans as blood hosts by *Ae. aegypti* mosquitoes (Rose et al., 2020; Rose et al., 2023). Furthermore, this variation in host preference was related to population density, seasonality, and level of precipitation, such that preference for human hosts increases with human population density and highly seasonal rainfall (Rose et al., 2020). In combined-variables models with the inclusion of ancestry and human population density, differences in host preference influence the variability of activity among our lines only during the day. This is not surprising as the absence of the host type that the individual lines have a strong preference for can influence their levels of activity during their most active time *i.e.* the daytime. In addition, ancestry was important for day activity and day sleep (with a stronger effect on day activity), which is unsurprising as significant biological differences have been associated with *Aaa*-like ancestry compared to lineages with high *Aaf*-like ancestry (Rose et al., 2020; Rose et al., 2023). The strong relationship of ancestry with activity and sleep parameters suggests an underlying genetic component of activity and sleep in mosquitoes, such as has been observed in *Drosophila* (Harbison et al., 2013). Establishing the specific genetic variants that influence activity and sleep in mosquitoes would require a more extensive genome-wide association analysis. The connection between ancestry and host preference might explain why we see similar effects concerning day activity for both explanatory variables. Moreover, human population density influences the differences in night activity and night sleep observed among our West African populations when the combined-variables models were considered. This implies the human availability for *Ae. aegypti* may have a considerable impact on differences in activity and sleep, especially during the night. This is consistent with earlier observations and predictions that host availability will modulate mosquito activity and/or sleep levels (Chaves et al., 2010; Rund et al., 2016; Ajayi et al., 2020). Increasing human population density and shifts to human preference, especially across sub-Saharan Africa, will cause mosquitoes to either modulate their activity and/or sleep cycle or duration in response to increased human availability and impacts of light pollution (Ajayi et al., 2020; Rund et al., 2020; Benoit and Vinauger, 2022; Ajayi et al., 2023). Due to urban expansion, the impact of artificial light at night (ALAN) on insect biology and biodiversity is increasing (Weng, 2014; Owens and Lewis, 2018). Apart from the known ecological effects of ALAN, such as the temporal disorientation of biological rhythms, especially in nocturnal insects, it is expected that continual exposure to light pollution would also have prolonged impacts on mosquito activity and sleep based on our results and previous ALAN studies (Rund et al., 2020), potentially leading to some lineages having altered sleep and activity levels as adaptations of prolonged residence in closer proximity to humans.

This research is the first study to examine intra-species differences in activity and sleep levels among *Ae. aegypti* lines and support earlier investigations in mosquitoes and other insects that activity and sleep traits are influenced by population or geographical differences even within the same animal species. Some similar observations were noted in *Ae. albopictus*, which revealed differences among populations for that species (Wynne et al., 2024). Future behavioral studies in mosquitoes will need to account for intra-species differences in activity and sleep that will be crucial to understanding disease transmission by mosquitoes, as this would improve current vector control strategies and predictive disease modeling.

## Acknowledgements and funding

This study was partially supported by the National Institute of Allergy and Infectious Diseases of the National Institutes of Health under Award Number R01AI148551 (to J.B.B. for shared incubator space), University of Cincinnati Sigma Xi (to O.M.A.), the National Institute of Food and Agriculture of the United States Department of Agriculture, Hatch project VA-1017860 and VA-160212 (to C.V.), and partially supported by the National Institute of Allergy and Infectious Diseases of the National Institutes of Health under Award number R01AI155785 (to C.V.). This study is directly supported by the National Institutes of Health R21AI166633 (to J.B.B. and C.V.), which focuses on understanding mosquito sleep. We thank Carolyn S. McBride for providing the mosquito populations used in this study.

## Supplemental materials

**Tables S1 - S16**: Estimated marginal means and pairwise post-hoc comparisons for all the lines. Each Excel worksheet represents data for the following parameters: total beam counts, day beam counts, night beam counts, total sleep amount, day sleep amount, night sleep amount, number of sleep bouts, and average sleep bout duration.

## Supplemental Figures

Figures S1 **– S17**: (A) Basic activity rhythms of 17 *Aedes aegypti* lines over a 24 h period. The y-axis represents the mean beam crosses in an activity monitor made by all the mosquitoes. (B) Sleep profiles of 17 *Ae. aegypti* lines averaged into a single 24 h period. The y-axis shows the proportion of time spent sleeping (defined as inactive periods of 120 min), averaged for each mosquito within a 30-minute time window. The x-axis for all the plots represents the Zeitgeber time (ZT0–ZT24). The solid lines and the shaded areas display means and their 95% bootstrap confidence interval, respectively. White and black horizontal bars represent the photophase and scotophase, respectively. KED: Kedougou (n=40); KIN: Kintampo (n=43); KUM: Kumasi (n=45); MIN: Mindin (n=41); NGO: Ngoye (n=36); PKT: PK10 (n=37); THI: Thies (n=34); COS: Costarica (n=40); D2M: D2MEB (n=37); GAI: Gainsville (n=52); IND: Indiana (n=48); LIV: Liverpool (n=36); BLA: Liverpool (Black eyes) (n=54); ORL: Orlando (n=48); PUI: Puerto-Ibo (n=41); PUR: Puerto Rico (n=37); ROC: Rockfeller (n=45).

## Notes

### Competing Interest Statement

The authors have declared no competing interest.

