## Supplemental Figures for "Intra-species quantification reveals differences in activity and sleep levels in the yellow fever mosquito, *Aedes aegypti*"

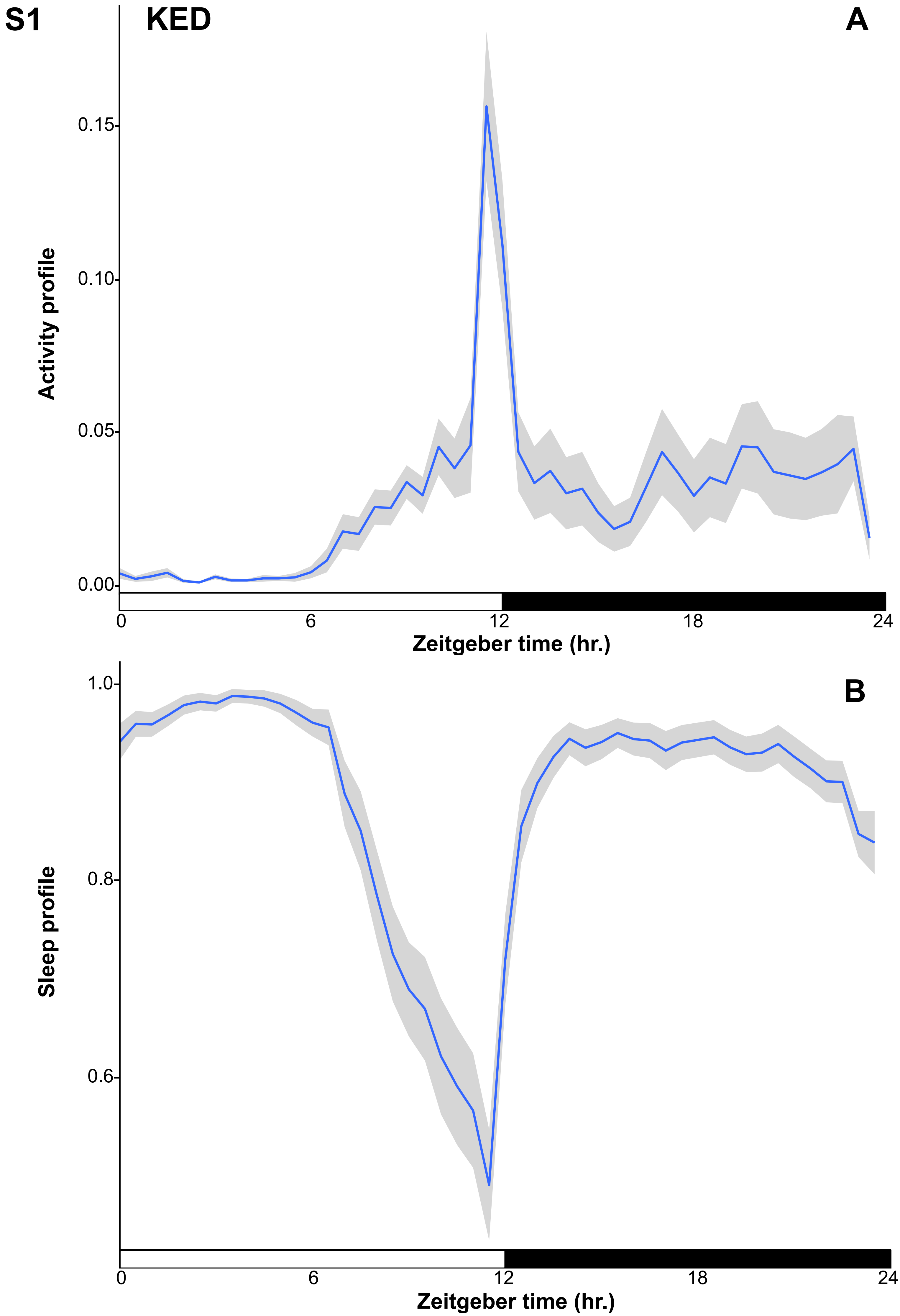

S1
KED
A
Activity profile
0.15
0.10
0.05
0.00
0
6
12
18
24
Zeitgeber time (hr.)
B
Sleep profile
1.0
0.8
0.6
0
6
12
18
24
Zeitgeber time (hr.)

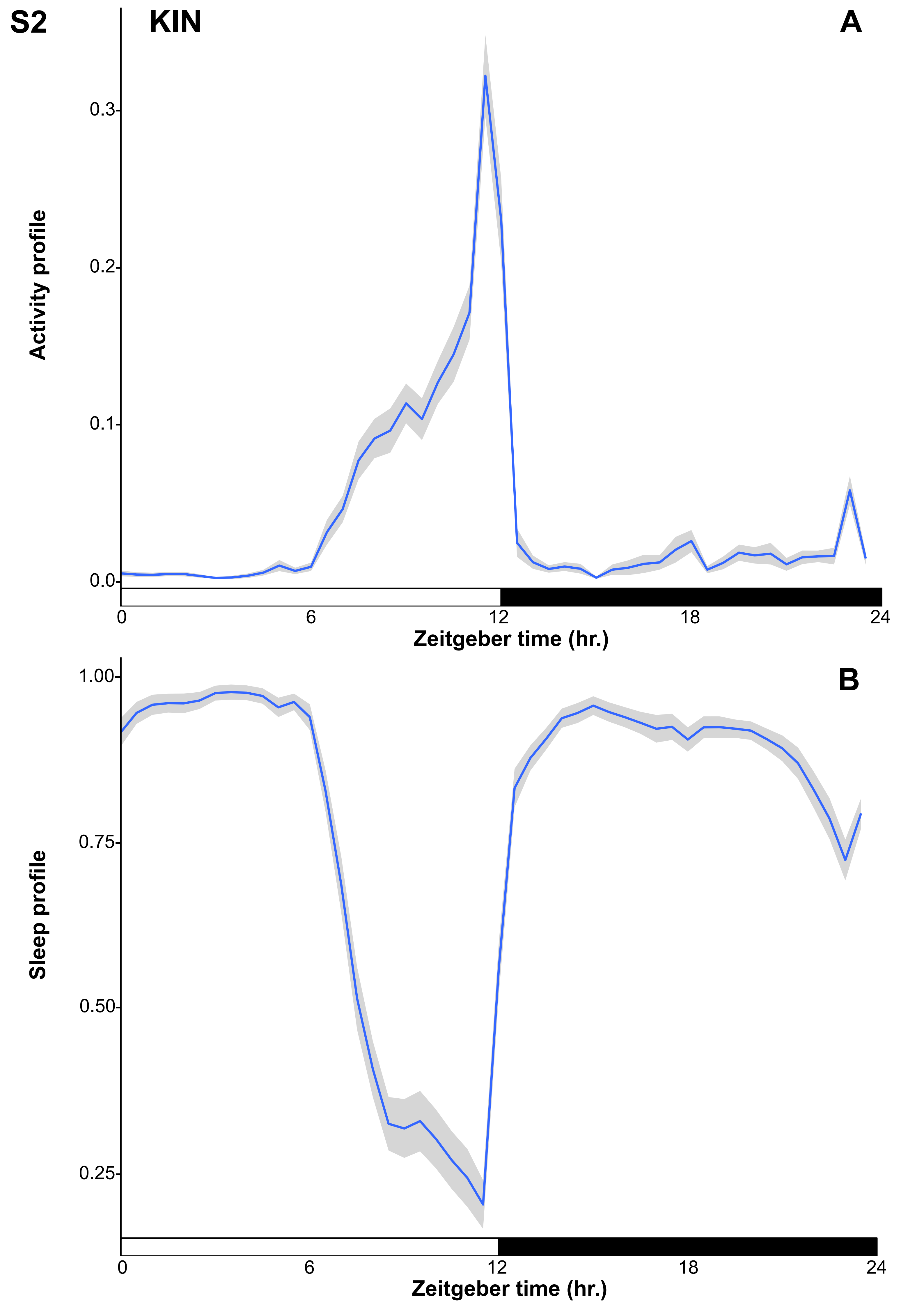
S2
KIN
A
Activity profile
0.3
0.2
0.1
0.0
0
6
12
18
24
Zeitgeber time (hr.)
B
Sleep profile
1.00
0.75
0.50
0.25
0
6
12
18
24
Zeitgeber time (hr.)

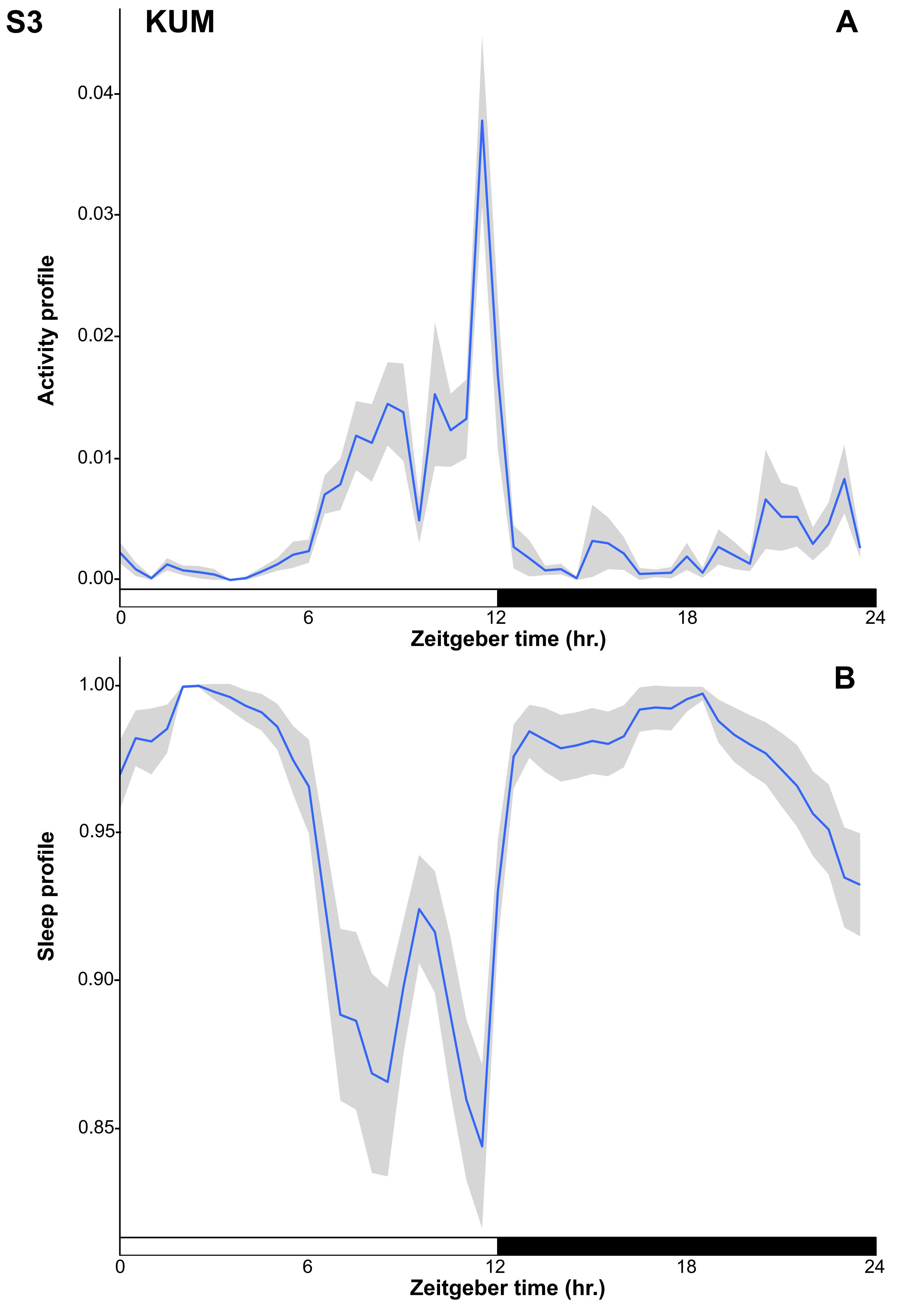

S3
KUM
A
Activity profile
0.04
0.03
0.02
0.01
0.00
0
6
12
18
24
Zeitgeber time (hr.)
B
Sleep profile
1.00
0.95
0.90
0.85
0
6
12
18
24
Zeitgeber time (hr.)

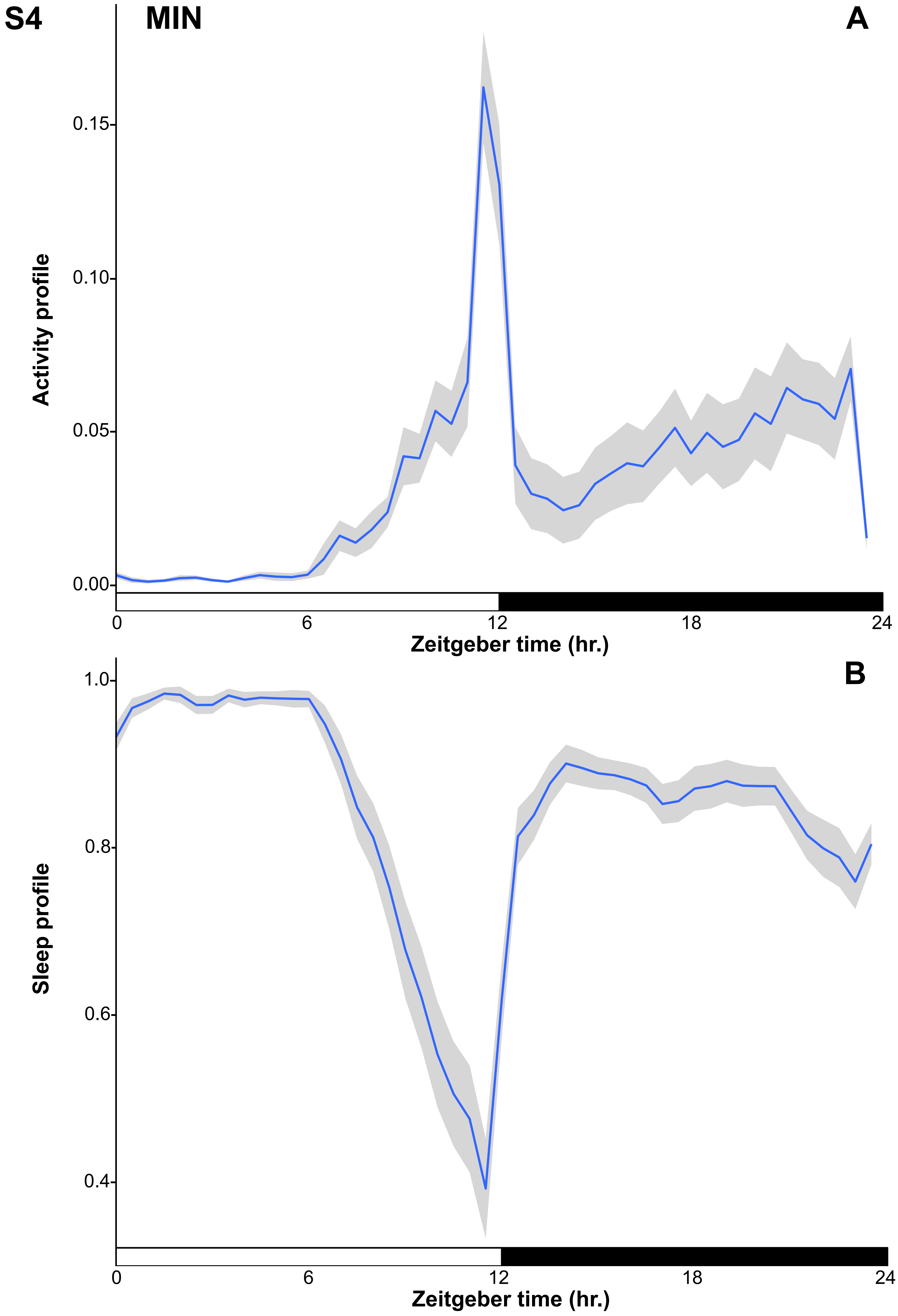

S4
MIN
A
Activity profile
0.15
0.10
0.05
0.00
0
6
12
18
24
Zeitgeber time (hr.)
B
Sleep profile
1.0
0.8
0.6
0.4
0
6
12
18
24
Zeitgeber time (hr.)

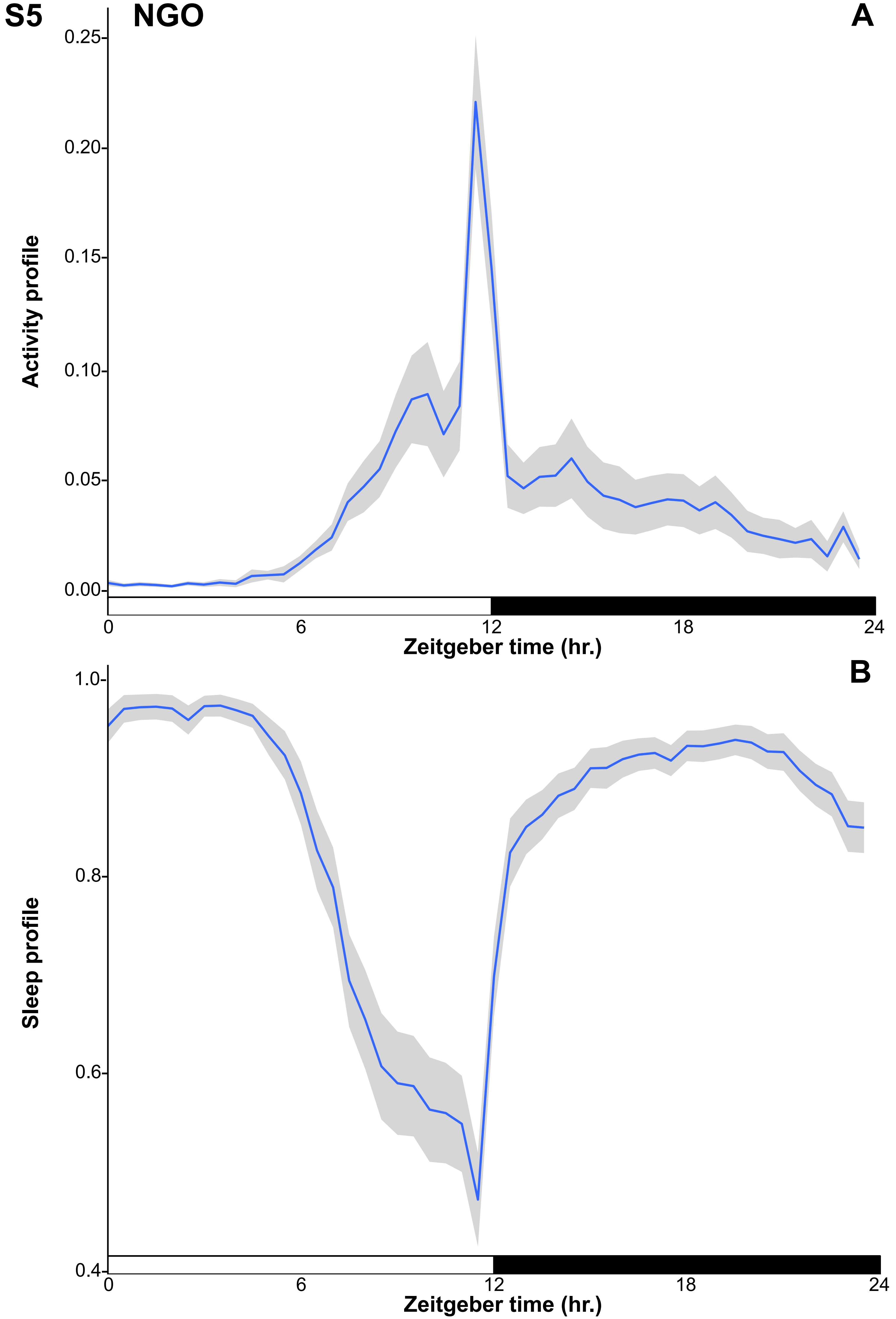
S5
NGO
A
Activity profile
0.25
0.20
0.15
0.10
0.05
0.00
0
6
12
18
24
Zeitgeber time (hr.)
B
Sleep profile
1.0
0.8
0.6
0.4
0
6
12
18
24
Zeitgeber time (hr.)

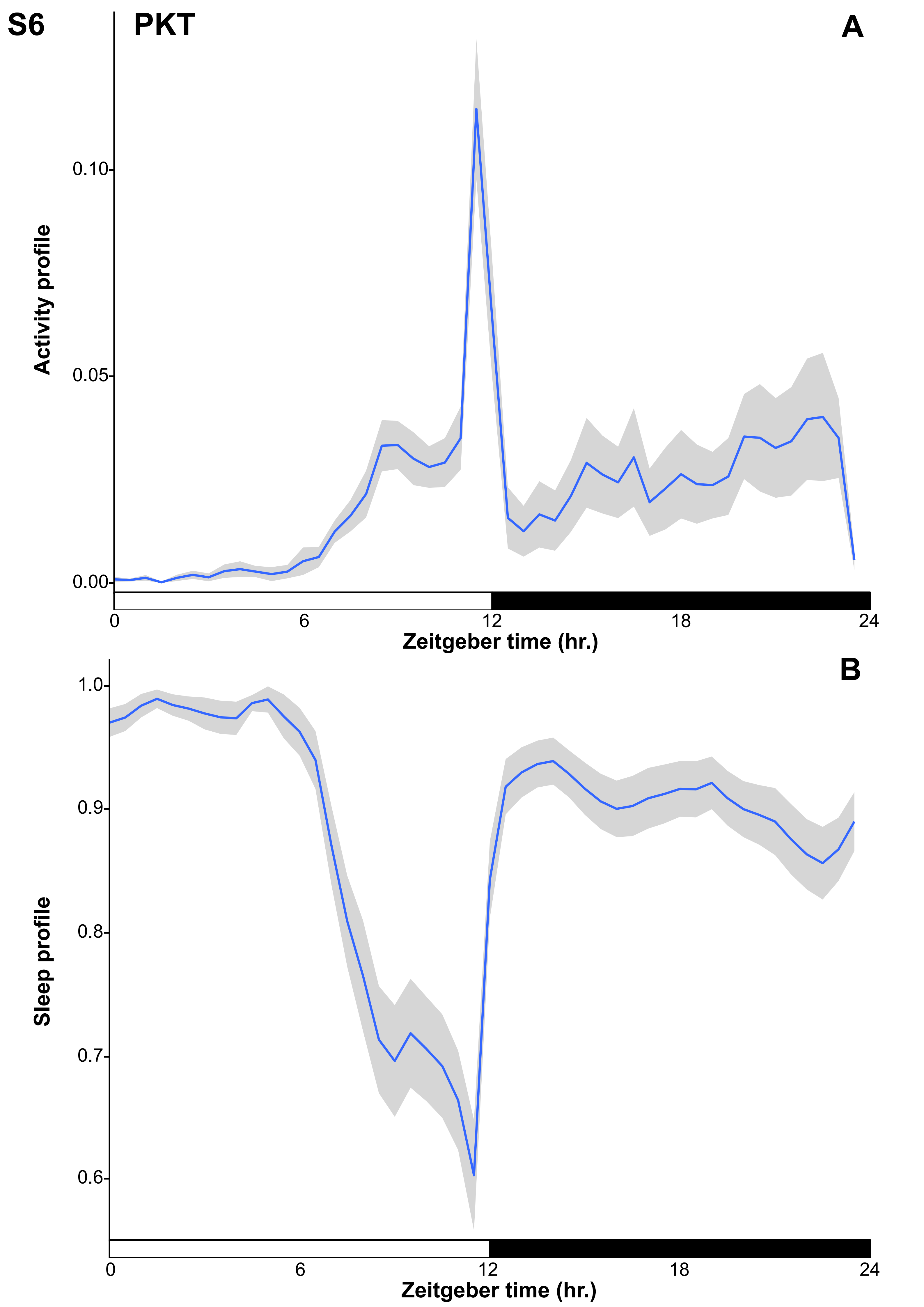

S6
PKT
A
Activity profile
0.10
0.05
0.00
0
6
12
18
24
Zeitgeber time (hr.)
B
Sleep profile
1.0
0.9
0.8
0.7
0.6
0
6
12
18
24
Zeitgeber time (hr.)

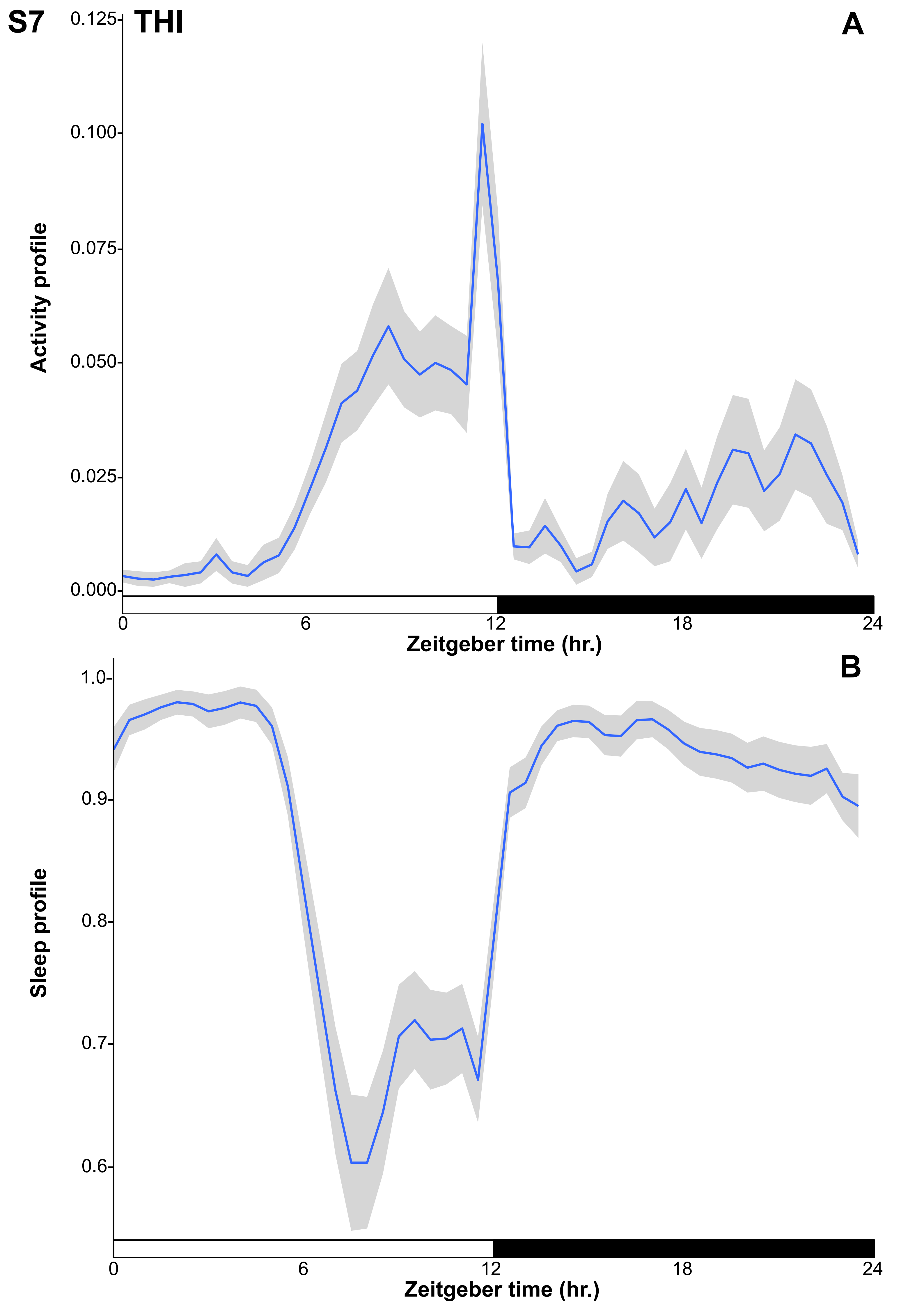
S7
THI
A
0.125
0.100
0.075
0.050
0.025
0.000
Activity profile
0
6
12
18
24
Zeitgeber time (hr.)
B
1.0
0.9
0.8
0.7
0.6
Sleep profile
0
6
12
18
24
Zeitgeber time (hr.)

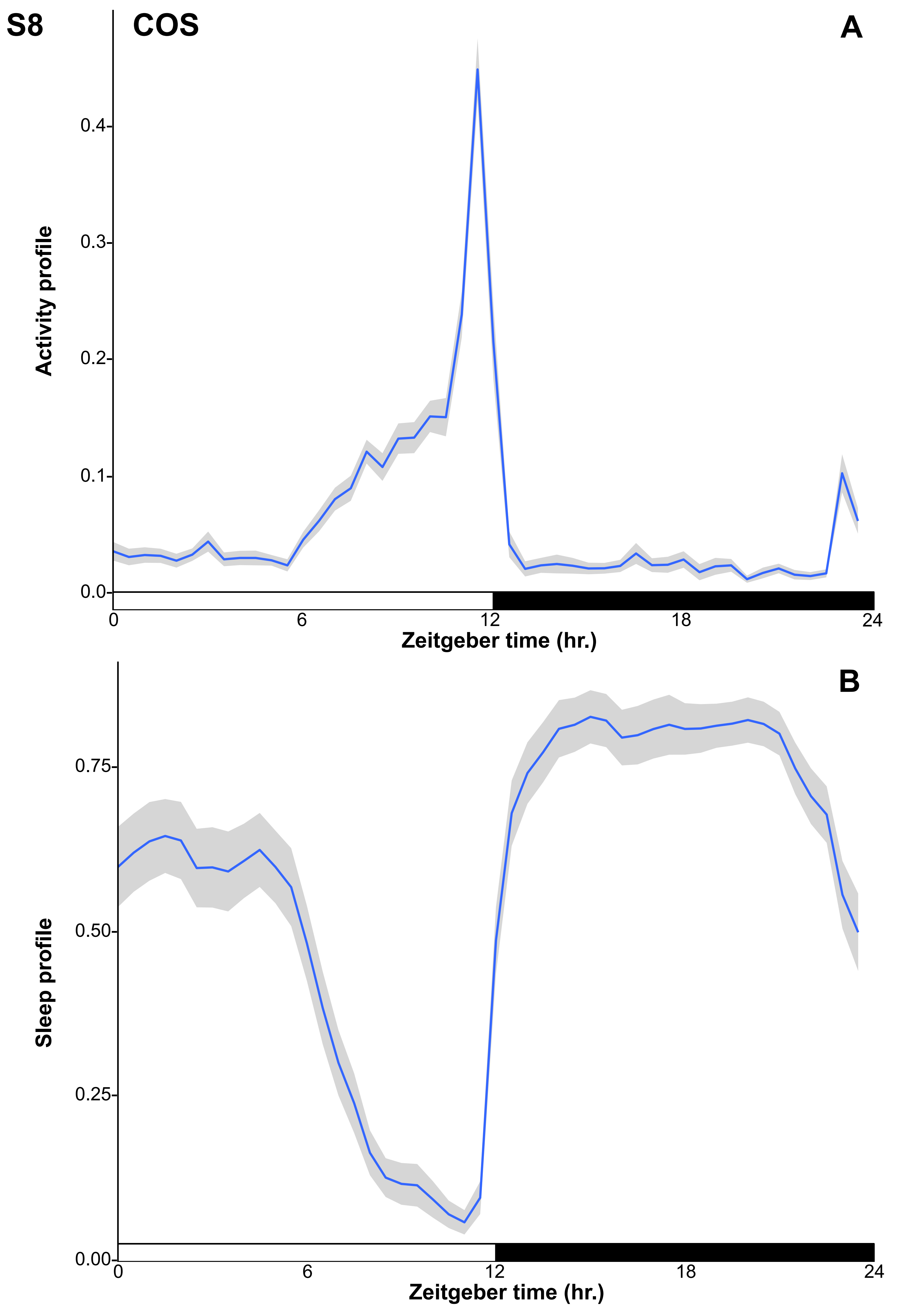
S8
COS
A
Activity profile
0.4
0.3
0.2
0.1
0.0
0
6
12
18
24
Zeitgeber time (hr.)
B
Sleep profile
0.75
0.50
0.25
0.00
0
6
12
18
24
Zeitgeber time (hr.)

S9

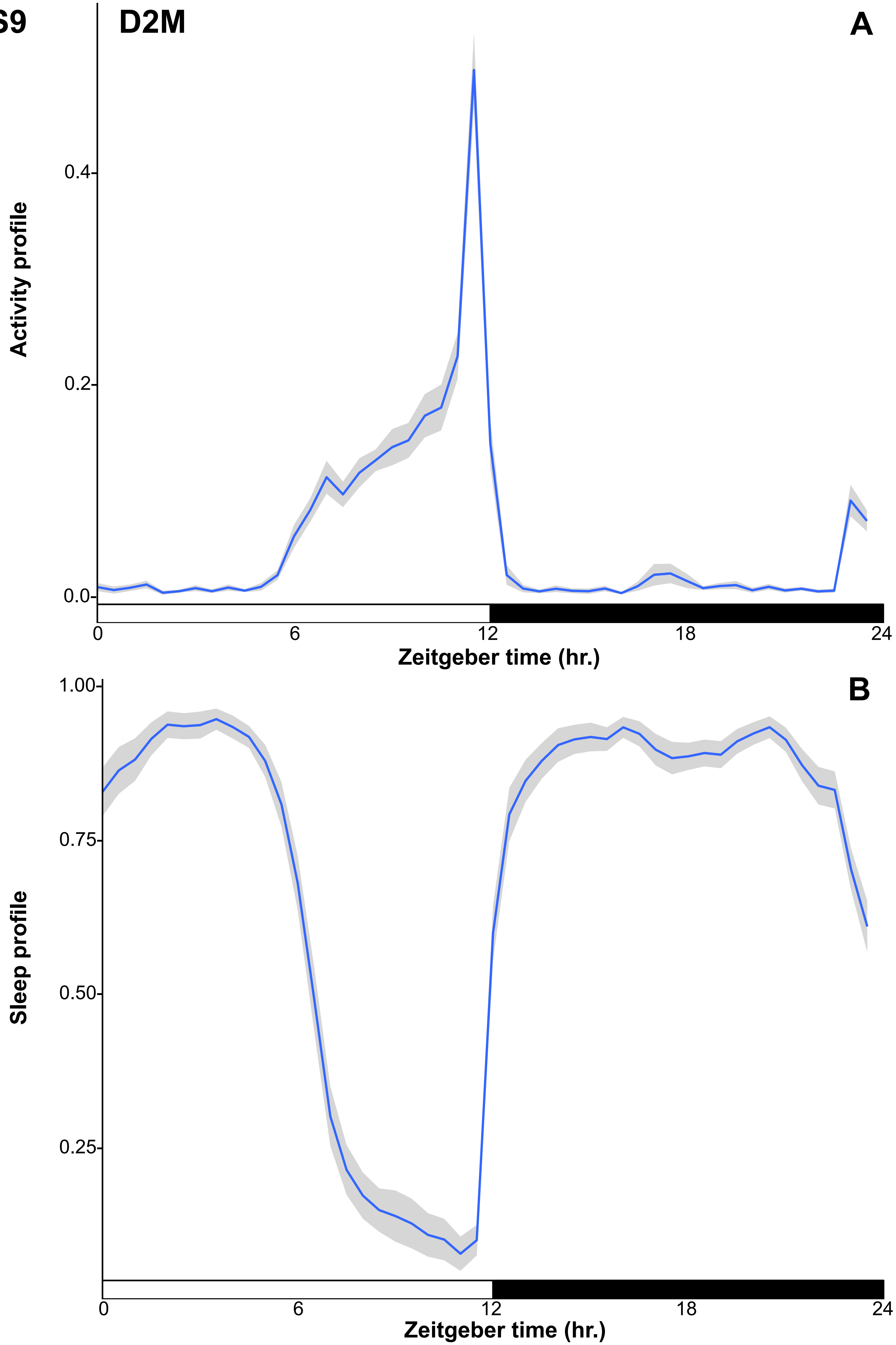

D2M
A
Activity profile
0.4
0.2
0.0
0
6
12
18
24
Zeitgeber time (hr.)
B
Sleep profile
1.00
0.75
0.50
0.25
0
6
12
18
24
Zeitgeber time (hr.)

S10

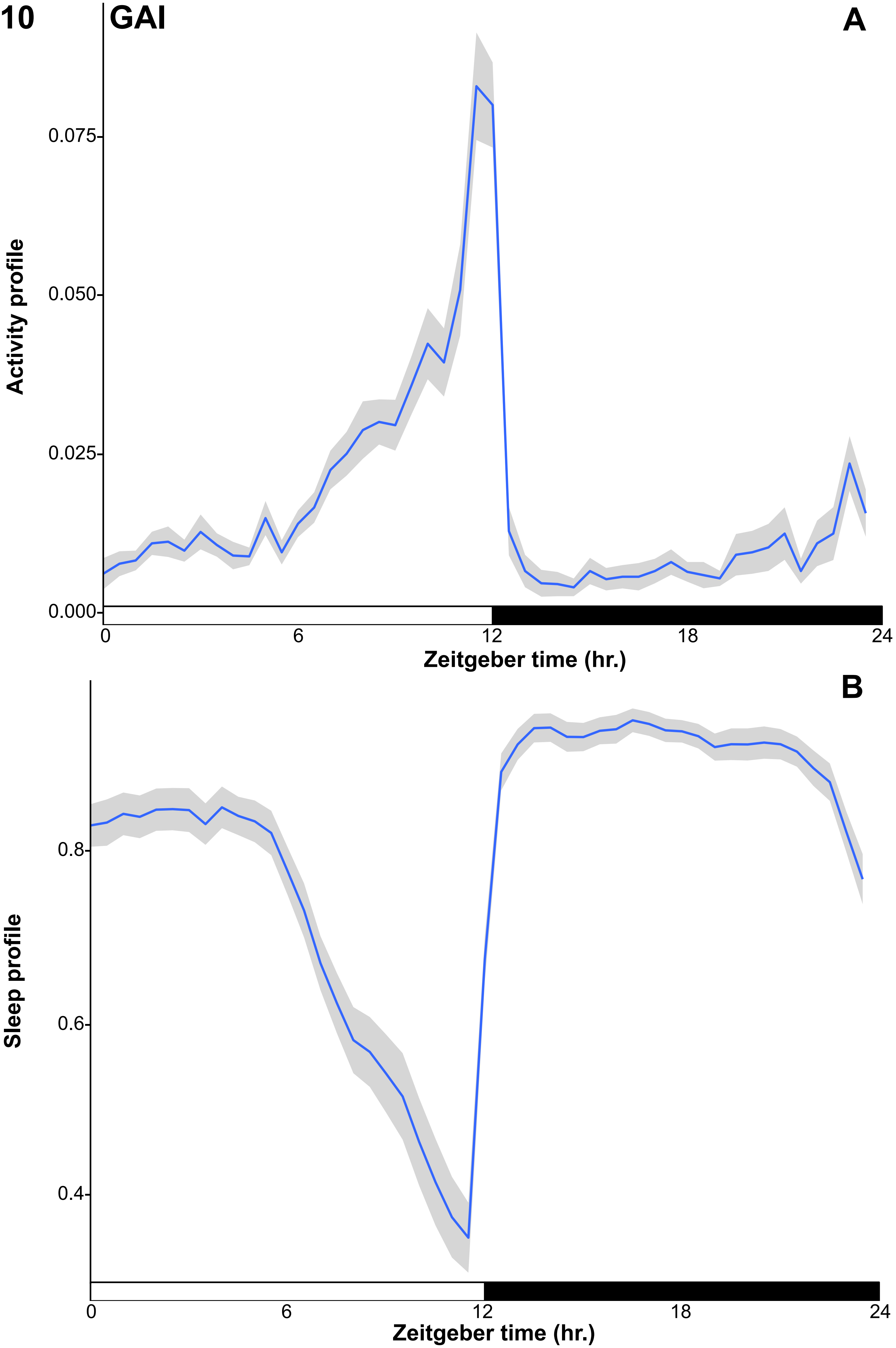
GAI
A
Activity profile
0.075
0.050
0.025
0.000
0
6
12
18
24
Zeitgeber time (hr.)
B
Sleep profile
0.8
0.6
0.4
0
6
12
18
24
Zeitgeber time (hr.)

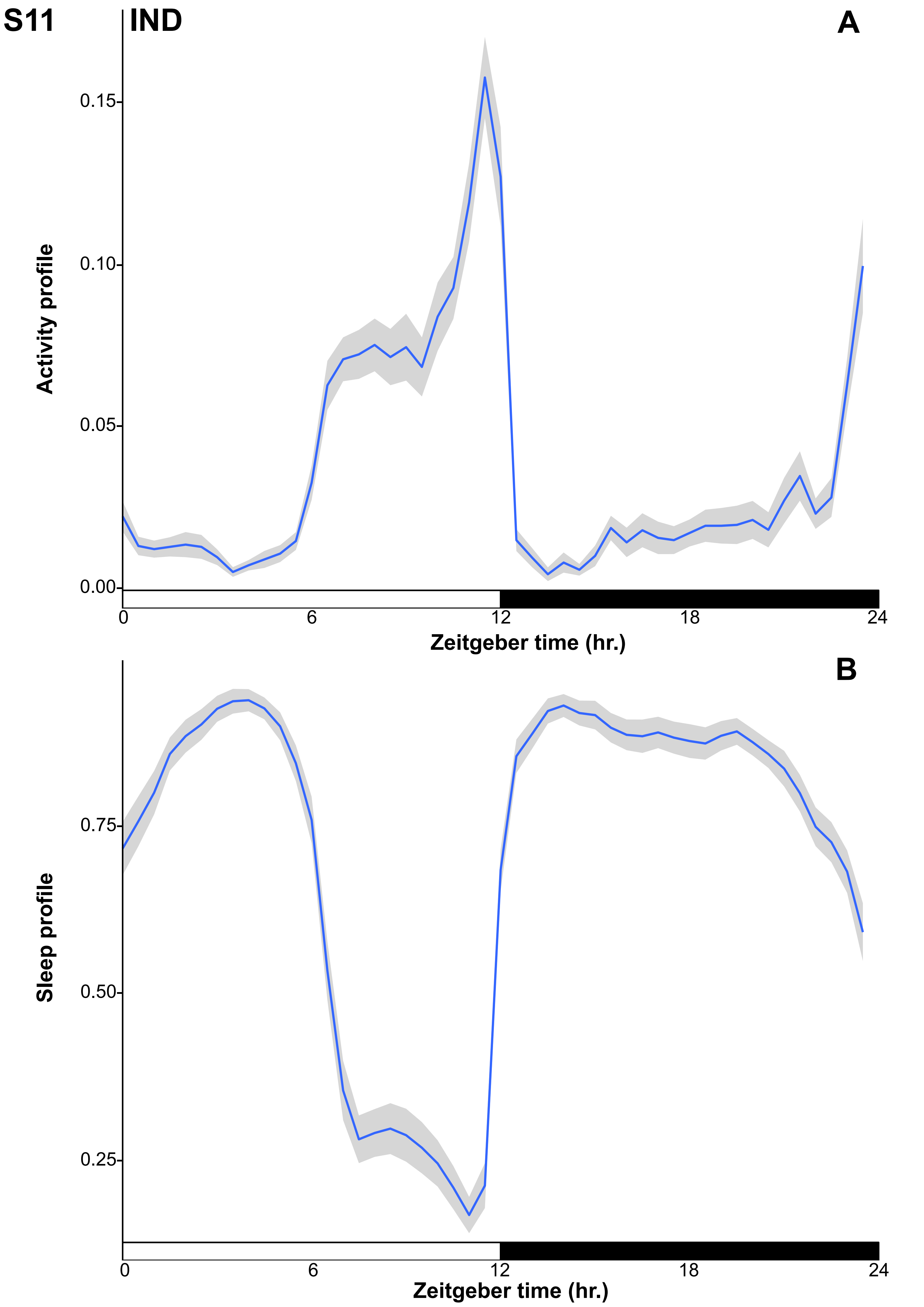
S11
IND
A
Activity profile
0.15
0.10
0.05
0.00
0
6
12
18
24
Zeitgeber time (hr.)
B
Sleep profile
0.75
0.50
0.25
0
6
12
18
24
Zeitgeber time (hr.)

S12

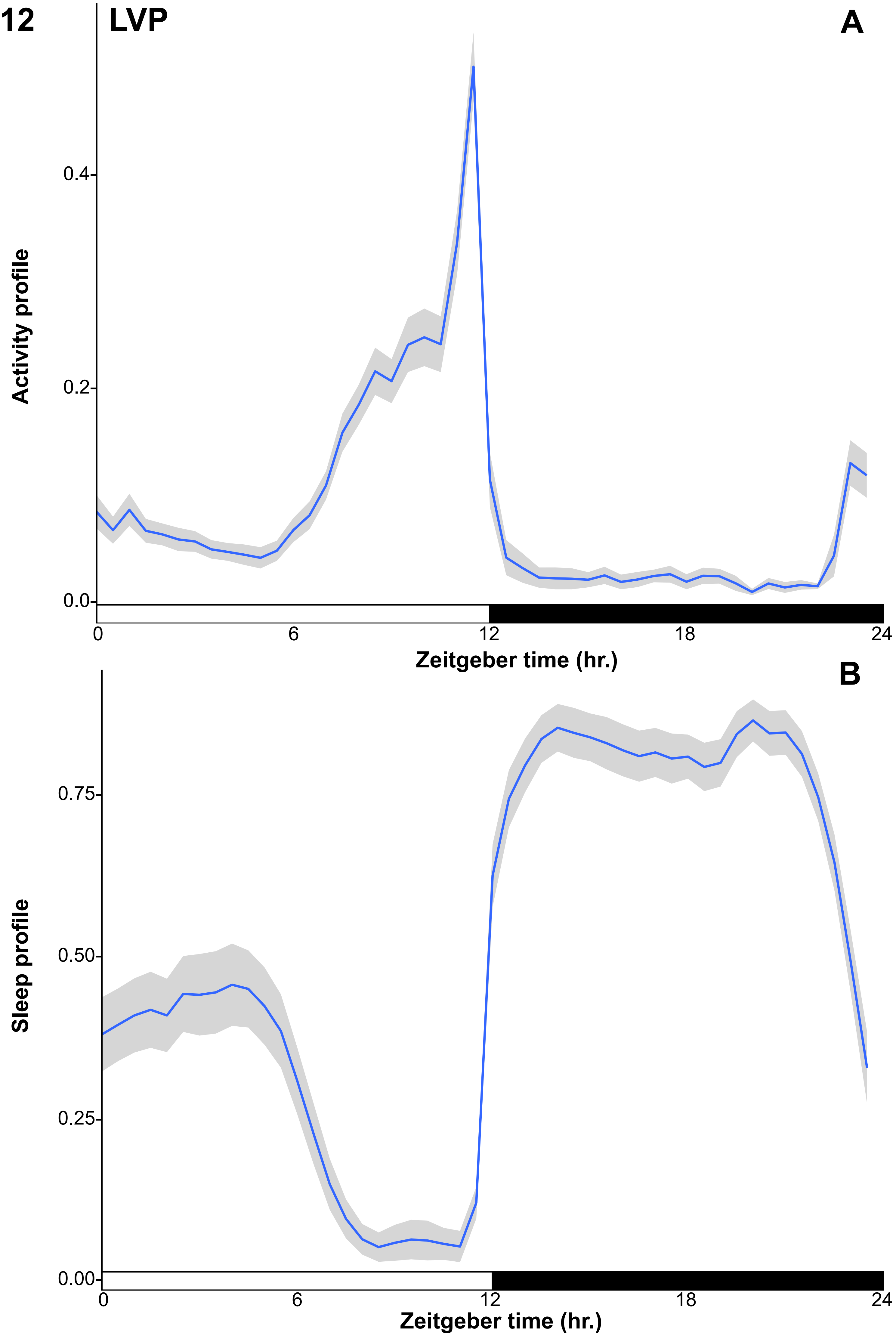
LVP
A
Activity profile
0.4
0.2
0.0
0
6
12
18
24
Zeitgeber time (hr.)
B
Sleep profile
0.75
0.50
0.25
0.00
0
6
12
18
24
Zeitgeber time (hr.)

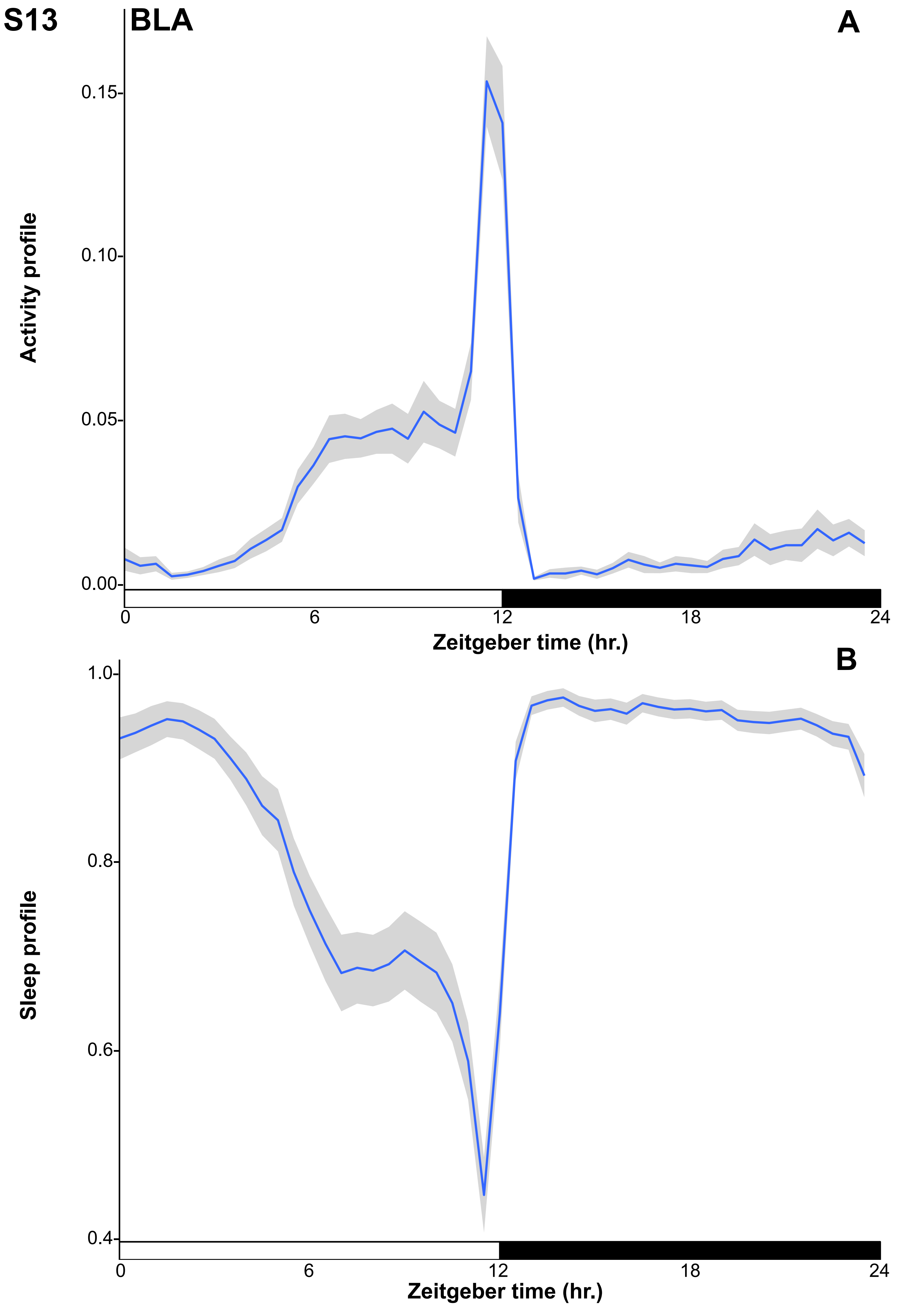
S13
BLA
A
Activity profile
0.15
0.10
0.05
0.00
0
6
12
18
24
Zeitgeber time (hr.)
B
Sleep profile
1.0
0.8
0.6
0.4
0
6
12
18
24
Zeitgeber time (hr.)

S14

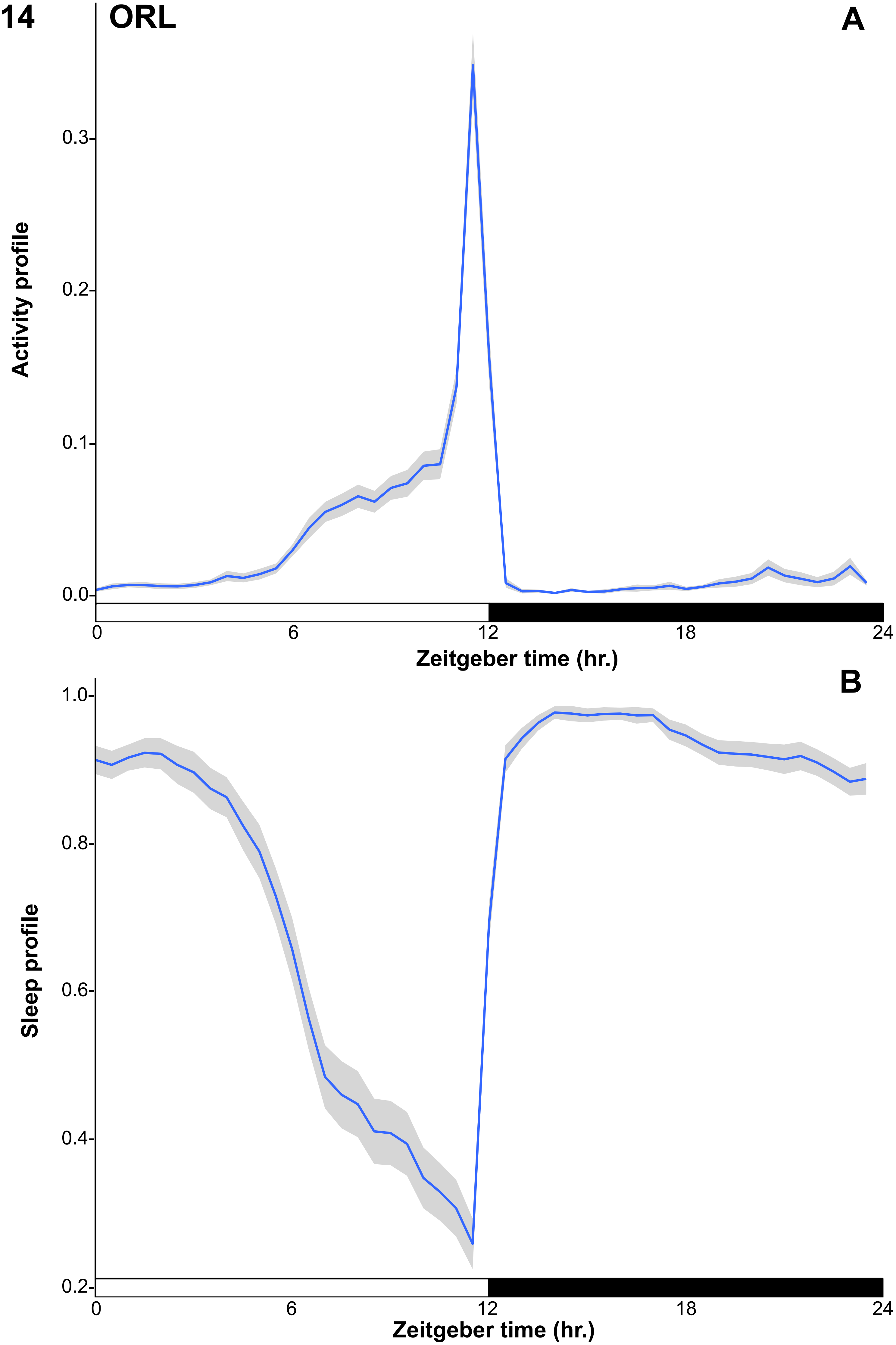
ORL
A
Activity profile
0.3
0.2
0.1
0.0
0
6
12
18
24
Zeitgeber time (hr.)
B
Sleep profile
1.0
0.8
0.6
0.4
0.2
0
6
12
18
24
Zeitgeber time (hr.)

S15

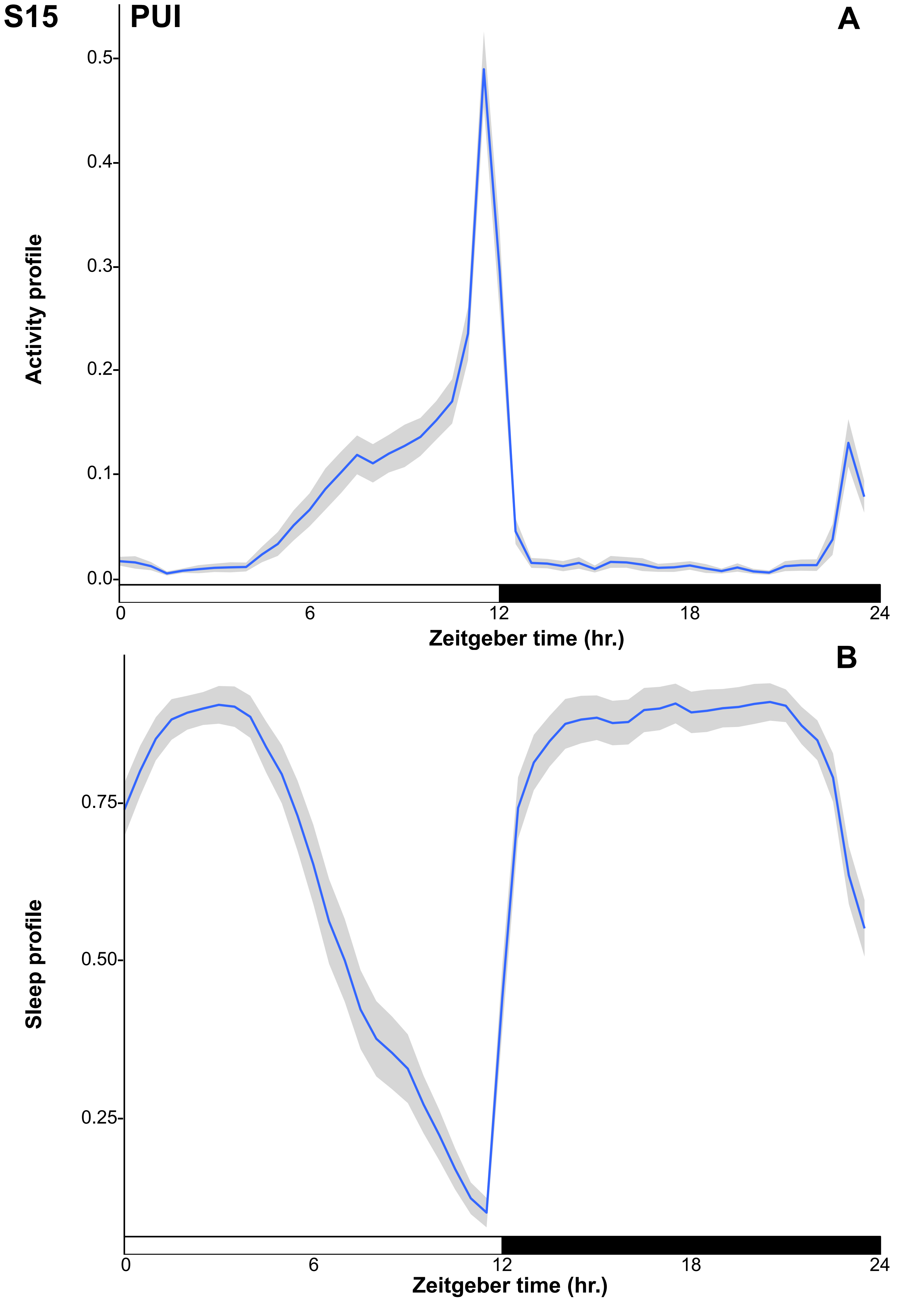
PUI
A
Activity profile
0.5
0.4
0.3
0.2
0.1
0.0
0
6
12
18
24
Zeitgeber time (hr.)
B
Sleep profile
0.75
0.50
0.25
0
6
12
18
24
Zeitgeber time (hr.)

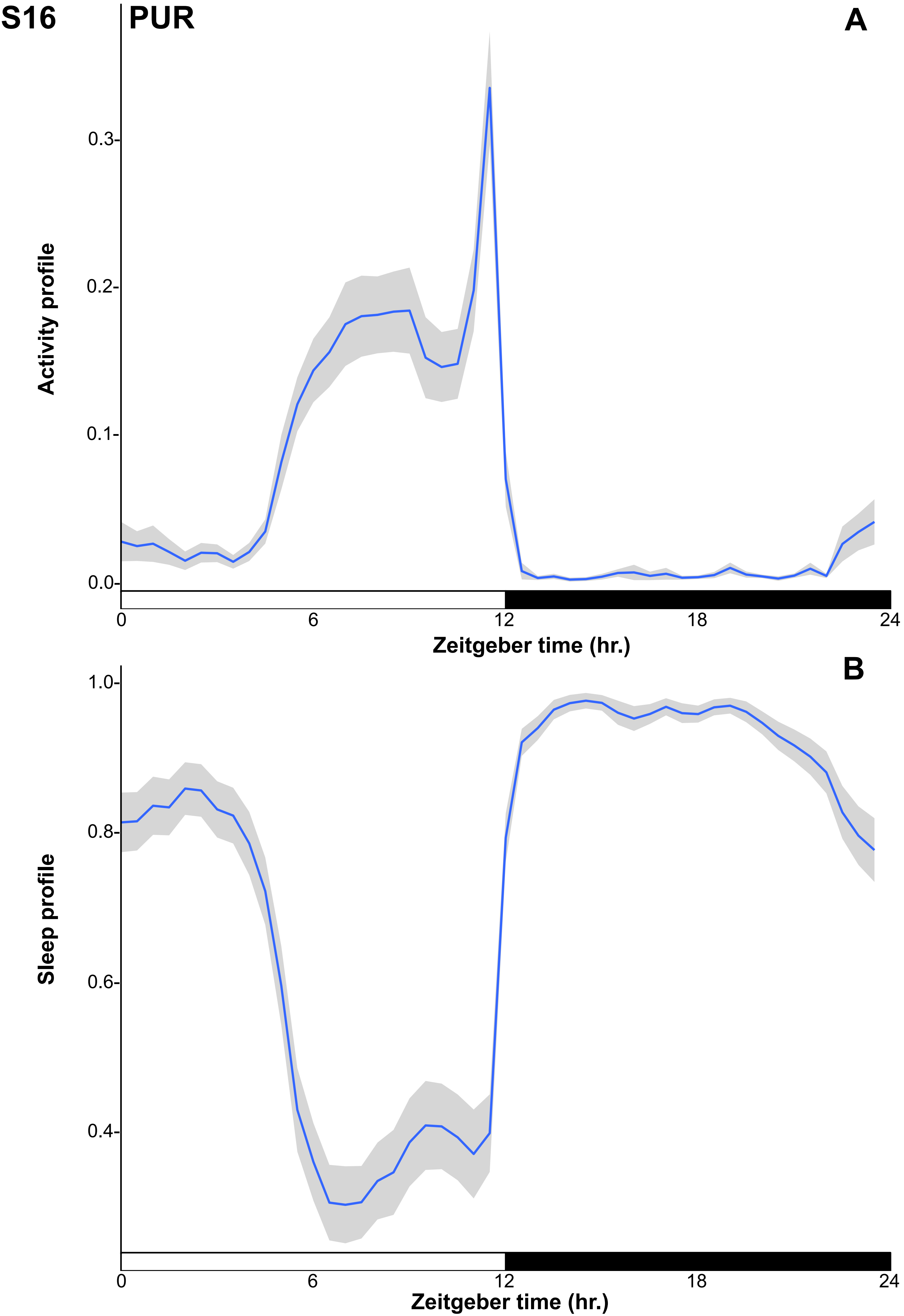
S16
PUR
A
Activity profile
0.0
0.1
0.2
0.3
0
6
12
18
24
Zeitgeber time (hr.)
B
Sleep profile
0.4
0.6
0.8
1.0
0
6
12
18
24
Zeitgeber time (hr.)

# S17

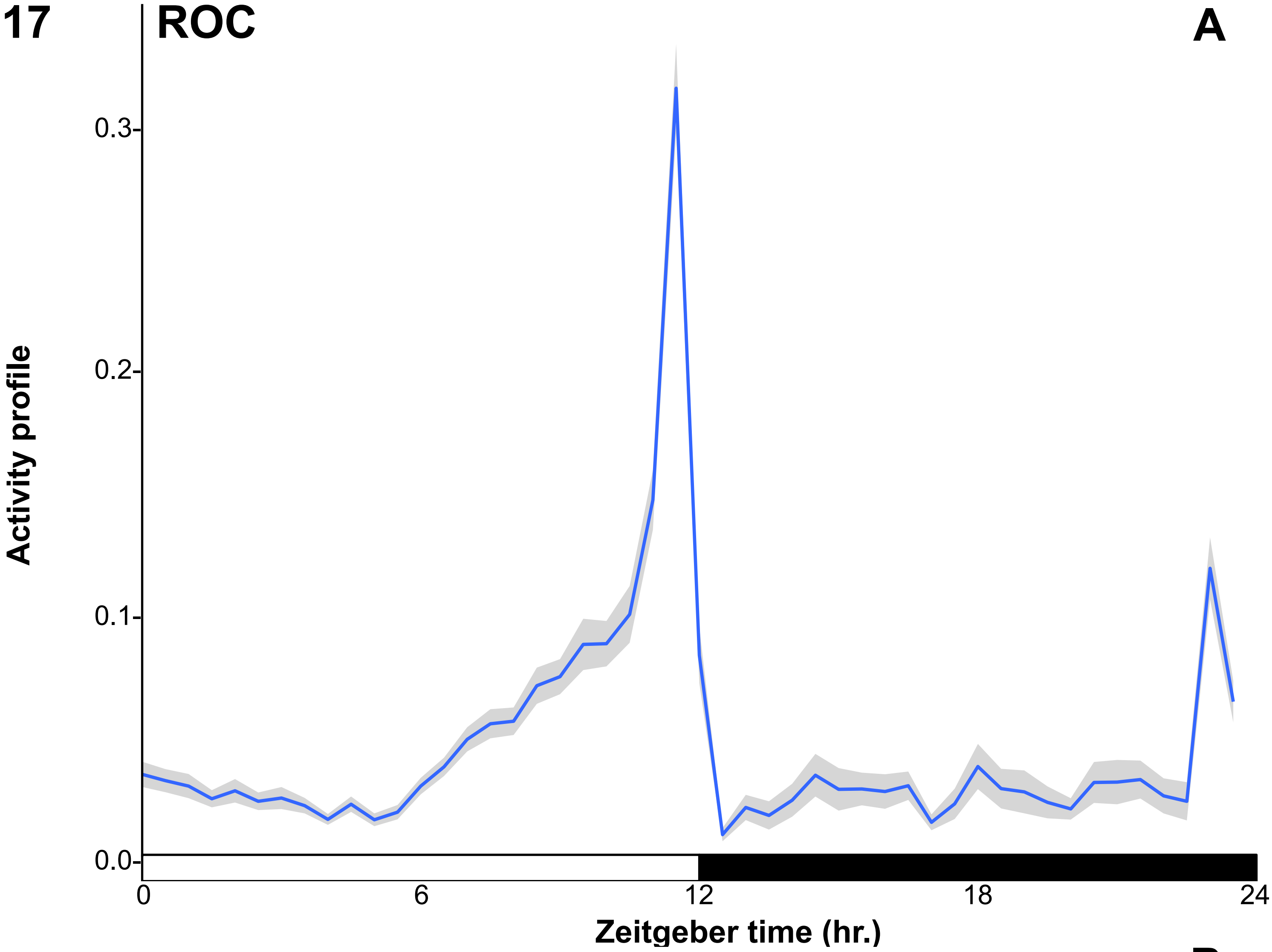
ROC
A
Activity profile
0.3
0.2
0.1
0.0
0
6
12
18
24
Zeitgeber time (hr.)

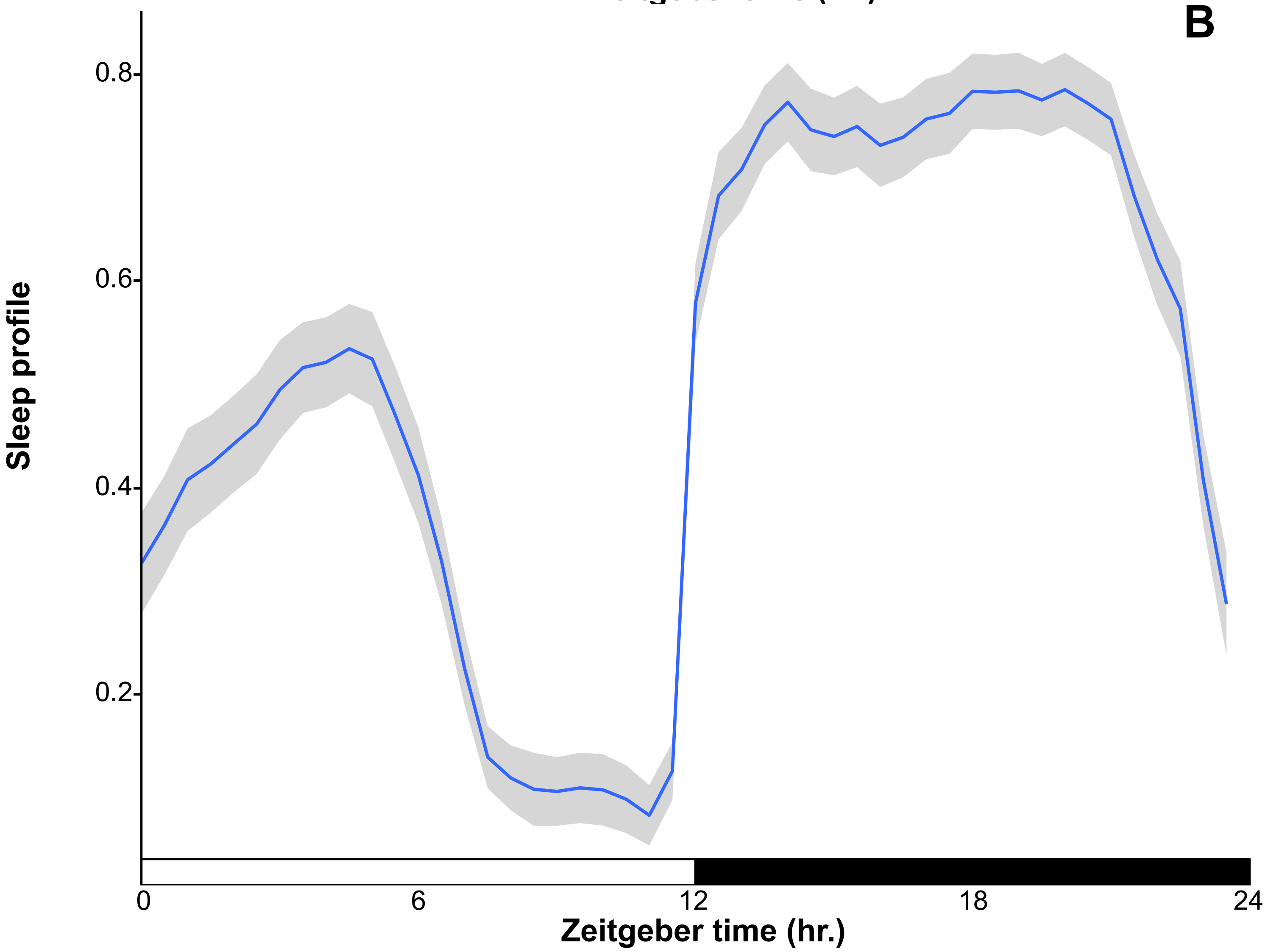
B
Sleep profile
0.8
0.6
0.4
0.2
0
6
12
18
24
Zeitgeber time (hr.)
